# Post-transcriptional regulation of *cyclin A* and *cyclin B* mRNAs is mediated by Bruno 1 and Cup, and further fine-tuned within P-bodies

**DOI:** 10.1101/2024.10.17.618951

**Authors:** Livia V. Bayer, Samantha N. Milano, Harpreet Kaur, Diana P. Bratu

## Abstract

Cell cycle progression is tightly controlled by the regulated synthesis and degradation of Cyclins, such as Cyclin A and Cyclin B, which activate CDK1 to trigger mitosis. Mutations affecting Cyclin regulation are often linked to tumorigenesis, making the study of cyclin mRNA regulation critical for identifying new cancer therapies. In this study, we demonstrate via super-resolution microscopy that *cyclin A* and *cyclin B* mRNAs associate with Bruno 1 and Cup in nurse cells. The depletion of either protein leads to abnormal Cyclin A and Cyclin B protein expression and a reduction in mRNA levels for both Cyclins. We further reveal that both *cyclin A* and *cyclin B* mRNAs accumulate in P-bodies marked by Me31B. Interestingly, Me31B is not involved in regulating *cyclin A* mRNA, as no changes in *cyclin A* mRNA levels or repression are observed upon Me31B depletion. However, *cyclin B* mRNA shows stage-specific derepression and reduced levels when Me31B is absent. Notably, the association between *cyclin B* and Cup is strengthened in the absence of Me31B, indicating that this interaction occurs independently of P-bodies. These results highlight the nuanced, mRNA-specific roles of P-body condensates in post-transcriptional regulation, challenging the idea of a uniform, binary mechanism of mRNA repression in P-bodies.

## Introduction

Under normal physiological conditions, cell cycle progression is tightly regulated by the controlled synthesis and degradation of mitotic cyclins. The cell cycle consists of four distinct phases: G0/G1 (Gap 1), S (DNA synthesis), G2 (Gap 2) and M (Mitosis). Each phase contains critical checkpoints to monitor DNA replication fidelity and chromosome segregation, ensuring proper cell division. Cyclins form complexes with cyclin-dependent kinases (CDKs), which, upon binding, activate the kinase activity of CDKs. These activated kinases subsequently phosphorylate key substrates that propel cell cycle progression. CDKs are highly stable proteins and exhibit consistent levels throughout the cell cycle. Temporal regulation of kinase activity is achieved via oscillation in the levels of their Cyclin binding partners (reviewed in [1]).

The activation of CDK1, the master regulator of mitosis, is sequentially mediated by Cyclin A (CycA), followed by Cyclin B (CycB), both of which are crucial for initiating and progressing through mitosis. During anaphase, CycB must be degraded by the anaphase-promoting complex/cyclosome (APC/C), an E3 ubiquitin ligase, to inactivate CDK1 and allow mitotic progression. Failure to degrade CycB results in a shortened G1 phase, abnormal DNA replication and genomic instability (reviewed in [2]). Notably, numerous cancers exhibit elevated levels of CycB, and a meta-analysis of 17 studies found that CycB overexpression correlates with poor survival outcomes in solid tumor patients. This makes CycB a promising therapeutic target, highlighting the need for further investigation into its regulatory mechanisms [3].

*Drosophila melanogaster* oogenesis serves as an excellent model system for studying cyclin mRNA regulation, as *cycA* and *cycB* mRNAs are post-transcriptionally regulated throughout the majority of oogenesis [4–8]. The unique expression pattern of these mRNAs enables the investigation of the specific mechanisms controlling mRNA regulation without the interference of the usual cyclical expression and degradation observed in other tissues. *Drosophila* oogenesis starts with the asymmetric division of a germline stem cell (GSC) in the germarium, producing a self-renewing stem cell and a differentiating cystoblast. The cystoblast undergoes four rounds of mitotic division with incomplete cytokinesis, forming a 16-cell cyst connected by cytoplasmic bridges known as ring canals. Of these 16 cells, 1 differentiates into the oocyte, while the remaining 15 develop into nurse cells. As the germline cyst develops, it becomes encapsulated by somatic follicle cells and exits the germarium as a fully formed egg chamber. This chamber subsequently undergoes 14 distinct developmental stages before being deposited into the oviduct (reviewed in [9, 10]).

After the egg chamber exits the germarium, only the oocyte nucleus remains in meiosis, arrested in prophase I, while the 15 nurse cells exit meiosis and enter endoreplication. During this process, the nurse cells replicate their DNA without undergoing cell division to supply the oocyte with the necessary factors for its growth. The proper regulation of GSC activity and the four rounds of mitotic division are controlled by the expression and degradation of Cyclins. Although the exact mechanism driving the cyst’s exit from the mitotic cycle remains unclear, the suppression of CycA and CycB protein expression is critical for this transition. CycA and CycB proteins are expressed in the germarium during the four rounds of mitotic divisions, and again starting at stage 12 of oogenesis. However, their mRNAs are expressed continuously throughout oogenesis and exhibit distinct localization patterns. *cycA* mRNA is evenly distributed throughout the egg chamber, whereas *cycB* mRNA preferentially accumulates in the oocyte until stage 7. After stage 7, the levels of both mRNAs decrease until a second round of transcription begins at stage 9, at which point these mRNAs are translated, setting the stage for proper embryonic development [4, 7, 8, 11].

The post-transcriptional regulation of *cycA* mRNA is mediated by Bruno1 (Bru1), an RNA-binding protein from the CELF1/2 family homolog in *Drosophila*, which directly binds to the 3’ UTR of *cycA* mRNA to repress its translation [7]. Bru1’s binding partner, Cup, an eIF4E-binding protein with a role in translational repression, also participates in the repression of *cycA* mRNA. In *bru1* and *cup* mutants, CycA protein is aberrantly expressed, leading to inappropriate mitotic re-entry [7]. This finding supports a model in which Bru1 binds to *cycA* mRNA’s 3’ UTR and recruits Cup, which in turn, binds to eIF4E to inhibit translation, resembling the proposed mechanism for *oskar* mRNA regulation [12]. CycB protein is also ectopically expressed in *bru1* and *cup* mutant egg chambers, but the regulatory mechanism involving Bru1 and Cup in CycB protein expression remains poorly understood. Notably, *cycB* mRNA lacks a predicted binding site for Bru1, which was found to only weakly bind to the 3’ UTR upon UV crosslinking [7]. Although the misregulated translation of *cycB* mRNA in *bru1* and *cup* mutants suggests both proteins have roles in its regulation, definitive evidence of them interacting with *cycB* mRNA has yet to be demonstrated.

Bru1 and Cup have been implicated in the regulation of numerous mRNAs and are considered core components of biomolecular condensates known as P-bodies (reviewed in [13]). P-bodies are cytoplasmic, membraneless organelles, formed via liquid-liquid phase separation (LLPS), where mRNAs can be stored in a translationally repressed state. Two key elements are essential for P-body formation: RNAs and RNA-binding proteins that contain intrinsically disordered regions. Once a local concentration threshold of these components is reached, LLPS is triggered [14–17]. The functional role of P-bodies remains a topic of ongoing debate. It remains unclear whether P-bodies form as a byproduct of high concentrations of mRNAs and associated proteins, leading to a thermodynamically favorable phase separation, or if they are actively involved in the mRNA life cycle, playing a crucial role in the spatio-temporal regulation of different transcripts (reviewed in [18]). This question highlights the need for further research to determine whether P-bodies are passive condensates or active regulators in mRNA processing. Nevertheless, it was found that approximately one-fifth of mRNAs localize to P-bodies as part of mRNA regulons. These mRNAs are translationally repressed and shielded from 5’ decay. Notably, the proteins encoded by these mRNAs are functionally related [19].

The localization of *cycA* and *cycB* mRNAs into P-bodies remains to be elucidated. *cycA* mRNA, which is bound by Bru1 and regulated by Cup, is a likely target for localization into P-bodies. *cycB* mRNA has been shown to localize to germ granules, another type of membraneless organelle that forms via LLPS [20]. Additionally, *cycB* mRNA undergoes poly(A) tail shortening via CCR4 and PABP2, followed by re-adenylation via Orb (CPEB protein) beginning at stage 10, prior to translation initiation [21]. A shortened poly(A) tail is a hallmark of mRNAs that localize into P-bodies (reviewed in [22]). Moreover, the CCR4-NOT complex, a major deadenylase in the egg chamber, interacts with Cup, which facilitates mRNA deadenylation while preventing subsequent decapping and degradation [15, 23–26]. Given these hallmarks, *cycB* mRNA is also a strong candidate for recruitment into P-bodies.

In this study, we confirm the association of Bru1 and Cup with *cycA* mRNA in the nurse cells, showing that the depletion of either protein results in aberrant CycA protein expression and a reduction in *cycA* mRNA levels. We show that *cycB* mRNA also associates with Bru1 and Cup in the nurse cell cytoplasm. Furthermore, downregulation of either Bru1 or Cup leads to rapid and robust CycB protein expression, accompanied by overall decreased mRNA levels in the egg chamber. Both *cycA* and *cycB* mRNAs accumulate in P-bodies, marked by the core P-body protein Me31B. We demonstrate that Me31B per se does not play a role in *cycA* mRNA translational regulation, as no ectopic expression of CycA protein was detected. The downregulation of Me31B does not affect the association of Cup with either mRNA, suggesting that the mRNP complex can form independent of P-body recruitment. On the contrary, stage-specific ectopic CycB protein was detected, suggesting that P-bodies play a supportive, fine-tuning role in regulating *cycB* mRNA.

## Results

### Bru1 and Cup knockdowns lead to re-entry into the cell cycle and developmental arrest

In our previous studies on the roles of Bru1 and Cup in *oskar* mRNA life cycle, we found that the depletion of both proteins causes developmental arrest [27]. We demonstrated that Bru1 and Cup regulate each other’s expression at the mRNA level, as the knockdown of either protein leads to a decrease in the levels of the other’s mRNA and protein. Furthermore, in *cup^RNAi^* egg chambers, endogenously tagged Bru1-GFP expression levels are decreased, and the cytoplasmic localization is disrupted. Bru1-GFP becomes diffuse throughout the cytoplasm and accumulates in large nuclear puncta (Figure 1A, and 1B, magenta arrowheads).

**Figure 1.**
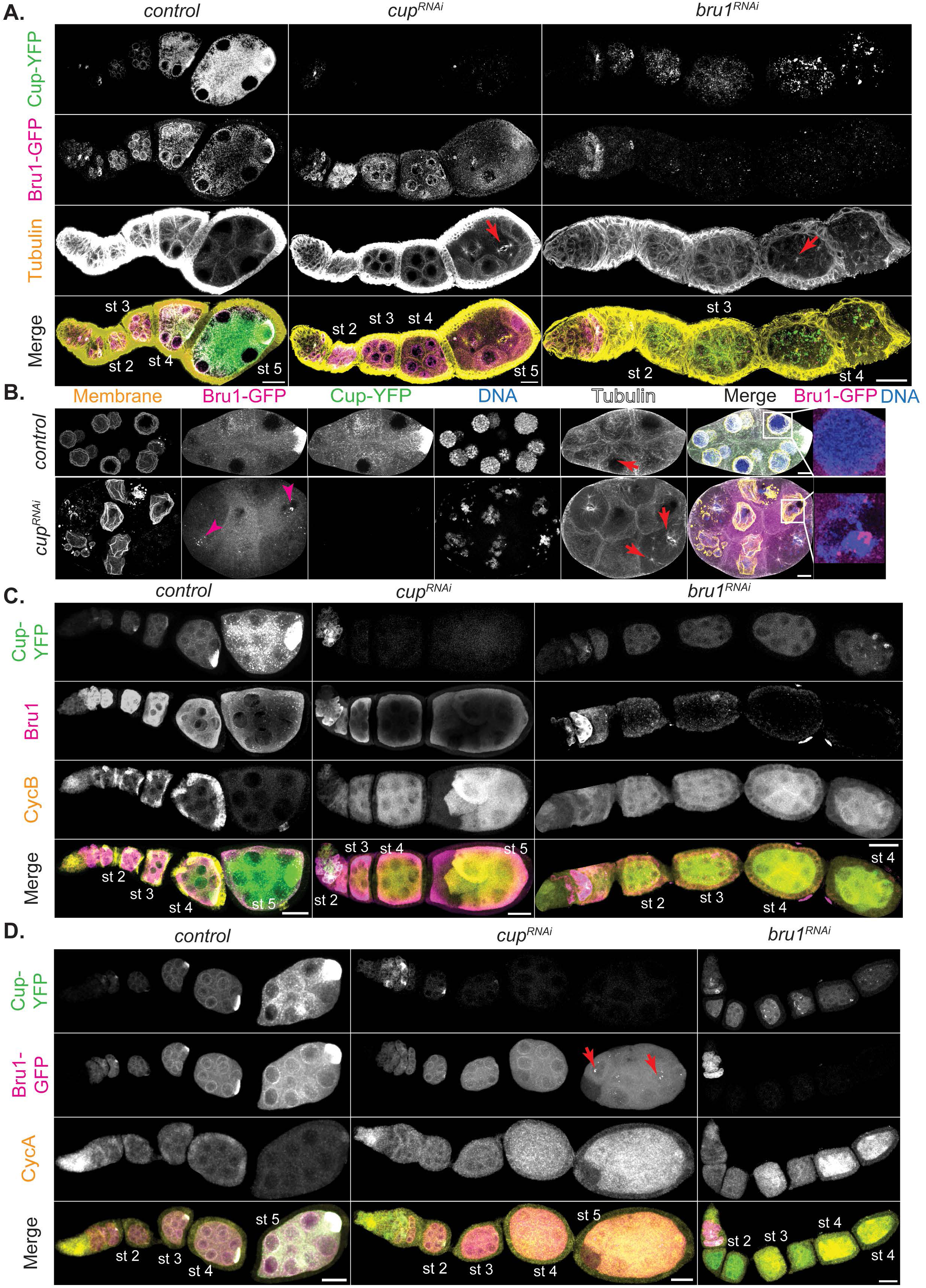
Bru1 and Cup knockdowns lead to re-entry of nurse cells into the cell cycle and to the ectopic expression of CycA and CycB proteins. (**A**) Visualization of –tubulin via IF in chains of egg chambers expressing Cup-YFP and Bru1-GFP. Formation of mitotic spindles in *cup^RNAi^* and *bru^RNAi^* backgrounds (red arrow). Images are deconvolved, XY max-intensity Z-projections of 5 optical slices (0.3µm each). Scale bars, 20μm. **(B)** Stage 6 egg chambers expressing Cup-YFP and Bru1-GFP in wild-type and *cup^RNAi^* backgrounds. Membrane (WGA stain). Nuclear aggregation of Bru1-GFP (magenta arrowheads and zoomed in merge panels). Formation of mitotic spindles (red arrows). Images are deconvolved, XY max-intensity Z-projections of 5 optical slices (0.3µm each). Scale bars, 10μm. **(C)** Visualization of CycB and Bru1 via IF in chains of egg chambers expressing Cup-YFP in the indicated backgrounds. Images are deconvolved, XY max-intensity Z-projections of 15 (*control*), 10 (*cup^RNAi^*) and 10 (*bru1^RNAi^*) optical slices (0.3µm each). Scale bars, 20μm. **(D)** Visualization of CycA via IF in chains of egg chambers expressing Cup-YFP and Bru1-GFP in the indicated backgrounds. Images are deconvolved, XY max-intensity Z-projections of 15 (*control*), 11 (*cup^RNAi^*) and 10 (*bru1^RNAi^*) optical slices (0.3µm each). Scale bars, 20μm.

We wished to more closely assess egg chamber development, using the UAS-Gal4 RNAi system to knock down Bru1 and Cup. We employed the Gal4 driver, *matalpha4-GAL-VP16*, in our studies to initiate knockdowns during early oogenesis (stages 1-2), thus allowing for Bru1 expression in the germarium, since the absence of Bru1 in the germarium causes a tumorous phenotype and blocks egg chamber formation. We found that Bru1 and Cup knockdowns distinctly affect progress through oogenesis (Figure 1A). Bru1 downregulation leads to a “beads on a string” phenotype, where multiple egg chambers in a single ovariole arrest during early stages of oogenesis, progressing just until stage 4. As they reach stage 4, the nurse cells re-enter the cell cycle, followed by the rapid degradation of the egg chambers (Figure 1A, red arrows; S1A).

Conversely, when Cup levels are reduced using the same driver, the egg chambers develop further, until stage 7 (Figure 1A). Yet, similarly to the Bru1 knockdown egg chambers, when limiting Cup levels, mitotic spindle formation is initiated at around stage 4 in multiple nurse cells as well as within multiple egg chambers in an ovariole (Figure 1B, S1B, red arrows). Unlike in the Bru1 knockdowns, once the nuclei divide, the nuclear envelope reforms, producing nuclei that are smaller in size. Whether this is due to multiple rounds of division, or to the reformation of the nuclei containing only partial chromosomal DNA remains to be determined (Figure 1B, S1B red arrowhead).

Altogether, these findings suggest that Bru1 plays a critical role in egg chamber maturation throughout early oogenesis, and not just in the germarium. Since Bru1 has been shown to repress the transcriptional repressor *polar granule component* (*pgc)* mRNA, it is possible that this derepression leads to global transcriptional changes that are detrimental to egg chamber development [28]. This activity seems independent of Cup, as *cup^RNAi^* egg chambers develop further than those with Bru1 depletion. These results demonstrate that both Bru1 and Cup are essential to suppress the re-entry of nurse cells into the cell cycle, possibly through the regulation of CycA and CycB proteins.

### CycA and CycB proteins accumulate at distinct developmental stages in Bru1 and Cup knockdown egg chambers

We aimed to visualize the ectopic accumulation of CycA and CycB proteins, as previous reports in *bru1* and *cup* mutant egg chambers showed the derepression of *cycA* and *cycB* mRNAs starting at the germarium stage [6, 7, 29]. To better understand their roles beyond the germarium, we focused on assessing ectopic CycA and CycB expression and mitotic re-entry in egg chambers where knockdowns were initiated post-germarium stages.

In wild-type egg chambers, CycA and CycB are expressed during the first four germ cell divisions in the germarium, and then only again after stage 12. Despite this expression gap, *cycA* and *cycB* mRNAs are continuously expressed throughout oogenesis [7, 30]. To investigate further, we knocked down Bru1 or Cup in egg chambers co-expressing endogenously tagged Bru1-GFP and Cup-YFP and assessed CycA and CycB distribution. Interestingly, we observed a robust, ectopic expression of CycB starting at stage 2 with either Cup or Bru1 depletion (Figure 1C, S1C), highlighting their critical roles in the translational repression of *cycB* mRNA. In contrast, CycA protein levels were low during early stages but increased starting at stage 4 in both knockdowns (Figure 1D, S1D). These patterns mirror the aberrant mitotic cycle initiation phenotype, with CycB alone being insufficient to trigger mitosis and CycA being required for the nurse cells to re-enter the cell cycle. This aligns with findings from earlier reports where CycB overexpression did not induce mitosis in nurse cells [31]. We confirmed the abnormal expression levels of CycA and CycB in these backgrounds via immunoblotting. We used 1-day old *mCherry^RNAi^* egg chambers as a control to limit the amount of late-stage egg chambers, where high levels of CycA and CycB are found (Figure S1E).

Surprisingly, in Bru1 knockdowns, CycA levels only significantly increase when Cup levels are severely reduced. Similarly, in Cup knockdowns, robust CycA expression is only detected when Bru1 levels in the cytoplasm decreases, and it accumulates in large nuclear puncta (Figure 1D, red arrows). The low CycA protein expression in earlier stages suggests that either Bru1 or Cup alone can partially repress *cycA* mRNA translation, but both are required for the strongest suppression. These results demonstrate that both Bru1 and Cup are essential for the translational repression of *cycA* and *cycB* mRNAs.

### Bru1 and Cup, with *cycA* and *cycB* mRNAs form large complexes in the nurse cell cytoplasm

Previous studies showed that *cycA* and *cycB* mRNAs are present throughout the egg chamber, with *cycB* mRNA accumulating in the oocyte. Additionally, although Bru1 directly binds *cycA* mRNA *in vitro*, it remains unclear whether it interacts with *cycB* mRNA, despite the ectopic accumulation of CycB protein in *bru1* mutant egg chambers [4, 7]. Similarly, the potential association between Cup and *cycA* or *cycB* mRNAs has not been explored. To fill these gaps, we aimed to 1) confirm the association between *cycA* mRNA and Bru1, 2) address whether Bru1 associates with *cycB* mRNA and 3) assess the interaction between *cycA* and *cycB* mRNAs with Cup in the nurse cell cytoplasm. Since highly disordered proteins, such as Bru1 and Cup, typically form complexes characterized by weak and highly dynamic interactions, co-immunoprecipitation experiments are often unreliable, particularly in *Drosophila* egg chambers. For example, in a highly sensitive study involving six RNA-binding proteins, numerous novel interactions were identified; however, some previously established interactions were not detected [32]. Additionally, the Egl/BicD/Dynein transport complex cannot be isolated through conventional immunoprecipitation techniques, although it can be successfully assembled *in vitro* [33, 34]. To overcome these limitations, we opted to visualize their association using sensitive single-molecule fluorescence *in situ* hybridization (smFISH) probes. Interestingly, we found that *cycA* mRNA, not only *cycB* mRNA, localizes into the oocyte, and that both mRNAs are present throughout oogenesis (Figure S2A).

To further assess *cycA* and *cycB* mRNAs’ localization and their spatial associations with Bru1 and Cup, we employed STED super-resolution microscopy in the nurse cell cytoplasm. As the large size of *Drosophila* egg chambers and their surrounding follicle cells present challenges for STED resolution, we developed a new protocol for sectioning egg chambers that proved to be essential for optimal STED imaging. We represent the zoomed-in images in a Punnett square format to display all possible combinations of colocalization (Figure 2A, S2B). We observed that both *cycA* and *cycB* mRNAs colocalize with Bru1-GFP and Cup predominantly within large cytoplasmic condensates (Figure 2A, S2B). We quantified these associations, and generated 2D plots for all our image data, where the x-axis represents the shortest distance between *cycA* or *cycB* mRNA particles to Bru1-GFP particles, and the y-axis reflects mRNA volume (Figure 2B, C). Our analysis reveals two distinct populations of mRNAs: a small subset, located further from Bru1, likely representing single mRNA copies, and a larger population closely associated with, or fully localized within, Bru1 condensates. The latter group, indicated by box plots below the x-axis, exhibits larger volumes, consistent with multiple mRNA copies (Figure 2B, C). Whether the single mRNA copies represent a subset that never interacts with Bru1, or are in transit to larger complexes, remains unclear. We propose that single-copy mRNAs are actively transported toward large cytoplasmic condensates, as small mRNA puncta are often found at the periphery of Bru1-GFP foci. A similar colocalization pattern is observed between Cup-YFP and *cycA* and *cycB* mRNAs (Figure 2A, D, E, S2C). Collectively, these results suggest that Bru1 and Cup associate with *cycA* and *cycB* mRNAs in the nurse cell cytoplasm, predominantly within larger cytoplasmic condensates.

**Figure 2.**
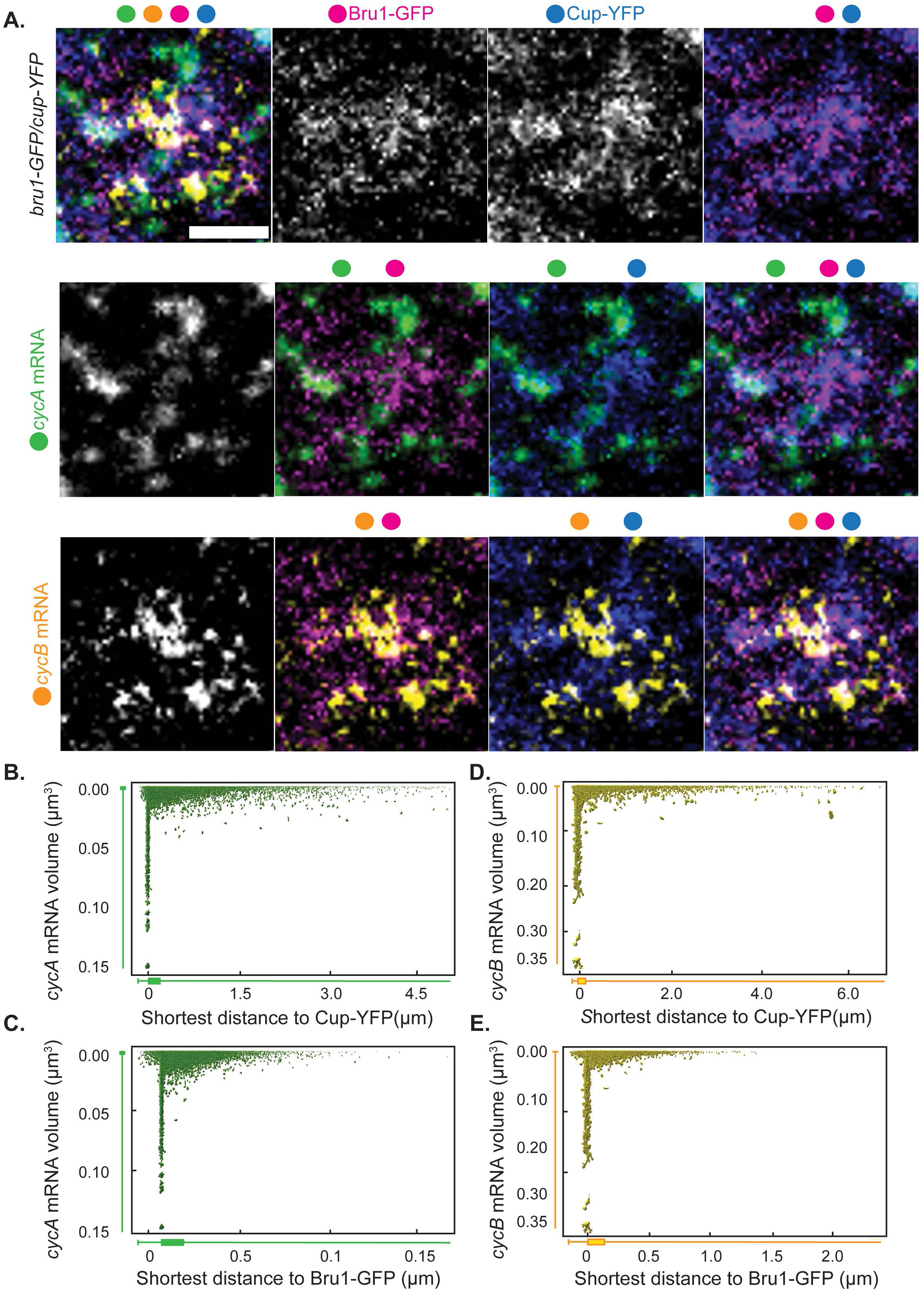
Bru1 and Cup form large complexes with *cycA* and *cycB* mRNAs in the nurse cell cytoplasm. **(A)** Co-visualization via STED of *cycA* and *cycB* mRNAs in nurse cells of a sectioned egg chamber at mid-oogenesis expressing Bru1-GFP and Cup-YFP. Images are laid out as a ‘punnett square’ for depicting colocalization combinations of RNAs and proteins. Colored dots indicate the individual channels merged per image. Images are deconvolved, XY max-intensity Z-projections of 3 optical slices (0.1µm each). Scale bar 1μm. **(B-E)** 2D-plots representing the distance to (µm – on the x-axis) and the volume of (µm^3^-on the y-axis) (**B**) Cup and *cycA* mRNA (**C**) Cup and *cycB* mRNA (**D**) Bru1 and *cycA* mRNA and (**E**) Bru1 and *cycB* mRNA.

### Bru1 and Cup are necessary for the formation of *cycA* and *cycB* mRNPs

Given that the knockdown of either Bru1 or Cup leads to ectopic expression of both CycA and CycB proteins, we next investigated the association of *cycA* and *cycB* mRNAs with either Bru1 or Cup in the absence of the other protein (Figure 3A). This allowed us to assess their respective roles in the formation of cytoplasmic mRNP complexes. We performed smFISH experiments, in combination with the RNAi-mediated knockdown of either protein. To ensure consistency, we quantified the colocalization of *cycA* and *cycB* mRNAs in egg chambers co-expressing Bru1-GFP and Cup-YFP at similar developmental stages, focusing on stages 2-4 (stage 4 was the oldest developmental stage observed in *bru1^RNAi^* egg chambers). We found that the knockdown of either Bru1 or Cup significantly diminished the association of the remaining protein with *cycA* and *cycB* mRNAs (Figure 3A). In the *bru1^RNAi^* egg chambers, the mean overlap ratio of the total particle volume of the *cycA* mRNA with Cup-YFP decreased by 74%, while that of *cycB* mRNA, it decreased by 58% (Figure 3B). Similarly, in *cup^RNAi^*, the mean overlap with Bru1-GFP was reduced by 70% for *cycA* and 63% for *cycB* mRNA, respectively (Figure 3C). Further analysis of stage 5-7 *cup^RNAi^* egg chambers revealed that Bru1-GFP and mRNA overlap was also severely reduced, with an 81% decrease for *cycA* mRNA and 74% for *cycB* mRNA (Figure S3A, B). Additionally, the overall mRNA levels for both the *cycA* and *cycB* transcripts were markedly reduced, as confirmed by RT-qPCR. Limited Cup expression led to a reduction of 80% in *cycA* mRNA and 40% in *cycB* mRNA levels (Figure 3D, E). The downregulation of Bru1 results in even more pronounced reductions, with 93% decreases in *cycA* mRNA and a 95% decrease in *cycB* mRNA levels (Figure 3D, E). These results demonstrate that both Bru1 and Cup are essential for establishing and/or maintaining the association between *cycA* and *cycB* mRNAs with the other component of the mRNP complex, as well as for preserving normal mRNA levels. Whether the observed reduction in mRNA levels was due to altered transcription, mRNA destabilization or degradation after translation remains to be discerned in future studies.

**Figure 3.**
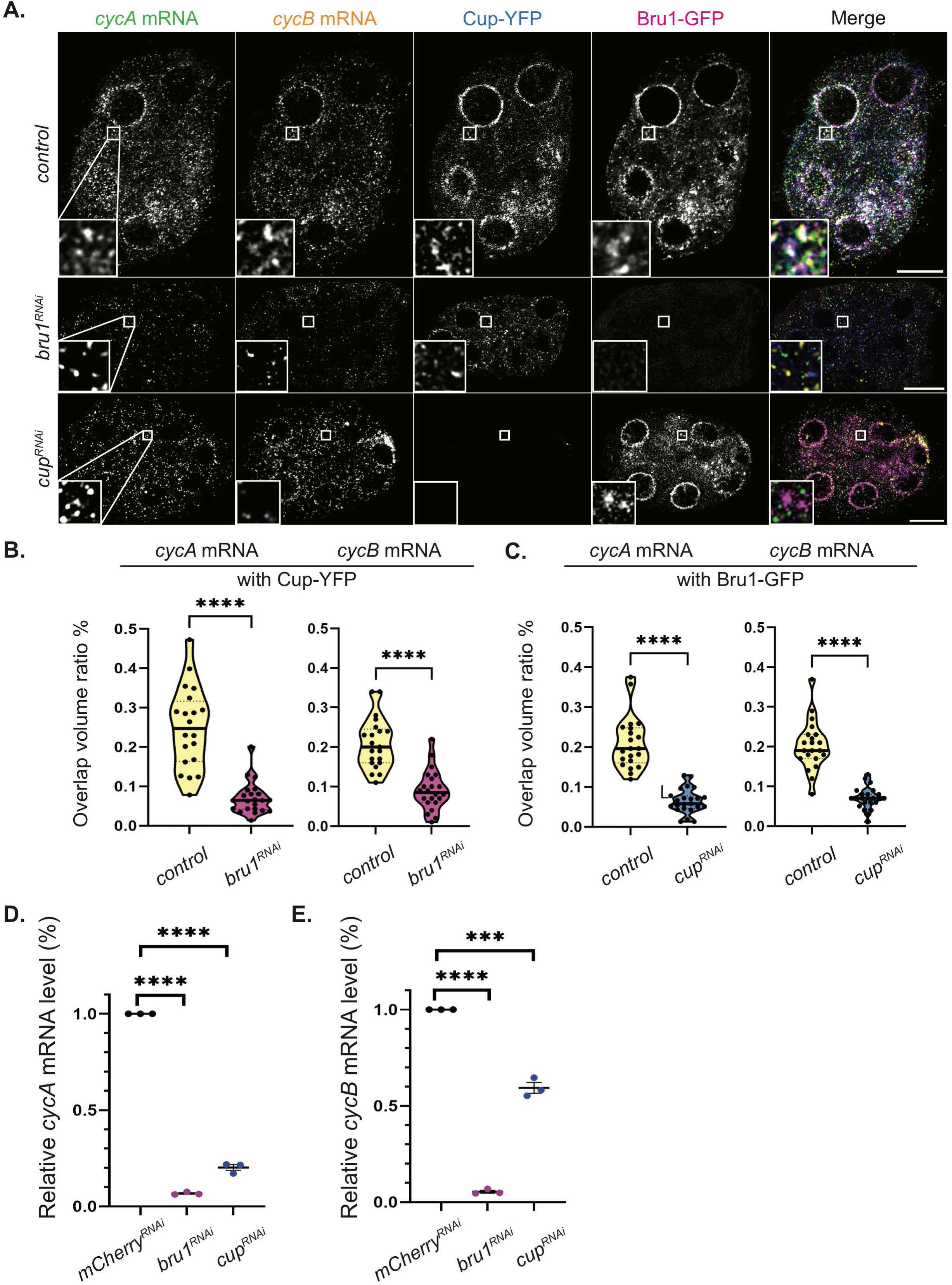
Bru1 and Cup are necessary for the formation of *cycA* and *cycB* mRNPs. **(A)** Co-visualization of *cycA* and *cycB* mRNAs in stage 4 egg chambers expressing Bru1-GFP and Cup-YFP in indicated backgrounds. White box indicates the location of the zoomed in images. Images are deconvolved, XY max-intensity Z-projections of 5 optical slices (0.3µm each). Scale bars, 10μm. **(B)** Overlap volume ratio analysis of *cycA* and *cycB* mRNAs with Cup-YFP performed in *control* and *bru1^RNAi^* stage 2-4 egg chambers (****p<0.0001; *control* vs *bru1^RNAi^*: n=20 and n=22) **(C)** Overlap volume ratio analysis of *cycA* and *cycB* mRNAs with Bru1-GFP performed in *control* and *cup1^RNAi^* stage 2-4 egg chambers (****p<0.0001; *control* vs *cup1^RNAi^*: n=21 and n=23) **(D)** RT-qPCR quantification of endogenous *cycA* mRNA normalized to *rp49* mRNA (mean ± SEM; ****p<0.0001; *mCherry^RNAi^* n=3, *cup^RNAi^* n=3). **(E)** RT-qPCR quantification of endogenous *cycB* mRNA normalized to *rp49* mRNA (mean ± SEM; ****p<0.0001; *mCherry^RNAi^* n=3, *cup^RNAi^* n=3).

### Re-entry into mitosis of Cup mutant egg chambers is rescued by Bru1 overexpression

In our analysis of knockdown egg chambers, mitosis is only observed at the beginning of stage 4 when both CycA and CycB are ectopically expressed, suggesting that the presence of both proteins is required for mitotic re-entry. *cycB* mRNA is rapidly derepressed, while *cycA* mRNA derepression only occurs when the levels of both Bru1 and Cup are significantly reduced. This raised the question of whether the overexpression of Bru1 or Cup could rescue the mitotic phenotype. To test this hypothesis, we generated fly lines to assess mitosis in a *cup* mutant background in combination with Bru1 overexpression.

In contrast to the *cup^RNAi^* egg chambers,the *cup* mutants developed further (∼ stage 8), with approximately 70% of ovarioles displaying at least one mitotic event. The overexpression of UAS-Bru1-GFP alone did not cause a phenotype at these stages when using the *matalpha4-GAL-VP16* driver. However, in the *cup^1/01355^* mutant background, UAS-Bru1-GFP overexpression significantly reduced the number of ovarioles with mitotic spindle formation to just 9% (Figure 4A, B). These egg chambers also progressed further in oogenesis, with most culminating in the “*cup*” phenotype. This differs from *cup* mutants, where most egg chambers die before reaching later stages, with only a few ovarioles displaying the “*cup*” morphology (Figure S4A).

**Figure 4.**
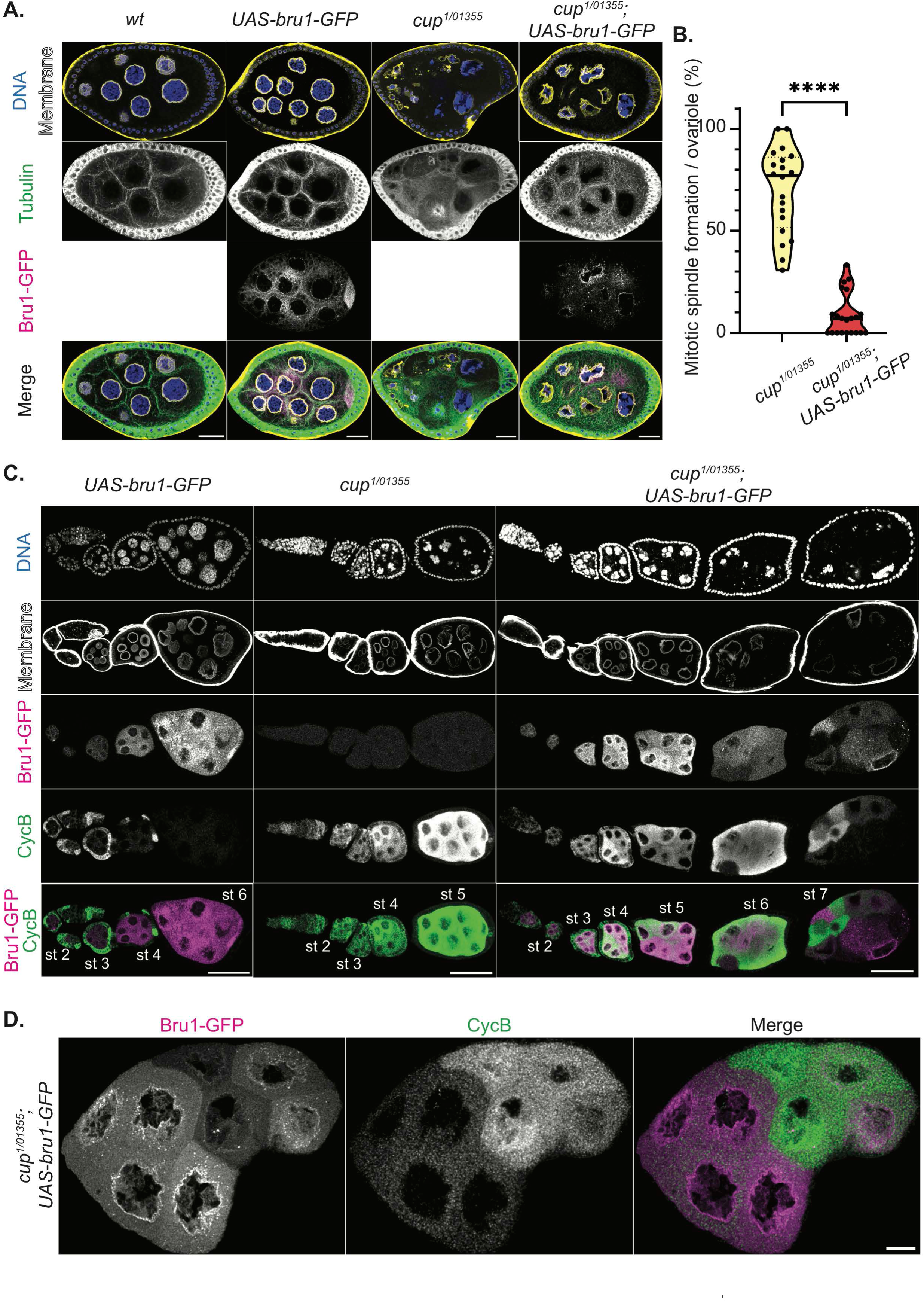
Bru1 overexpression rescues mitotic re-entry of *cup* mutant egg chambers. **(A)** Stage 6 egg chambers stained for –tubulin via IF in the indicated backgrounds. Membrane (WGA). Images are deconvolved, XY max-intensity Z-projections of 3 (*wt*), 3 (*UAS-bru1-GFP*), 3 (*cup^1/01355^*) and 4 (*cup^1/01355^; UAS-bru1-GFP*) optical slices (0.3µm each). Scale bars, 20μm. **(B)** Analysis of mitotic spindle formation in *cup^1/01355^* mutant egg chambers and in *cup^1/01355^* mutant egg chambers expressing the UAS-Bru1-GFP transgene. Each data point represents the percentage of ovarioles containing at least one mitotic spindle formation event in *cup^1/01355^*(n=20) and *cup^1/01355^*; *UAS-bru1-GFP* (n=21) ovaries. ****p<0.0001 **(C)** Chains of egg chambers in the indicated backgrounds, stained for CycB via IF. Membrane (WGA). Images are deconvolved, XY max-intensity Z-projections of 5 (*UAS-bru1-GFP*), 6 (*cup^1/01355^*) and 6 (*cup^1/01355^; UAS-bru1-GFP*) optical slices (0.3µm each). Scale bars, 50μm. **(D)** Stage 7, *cup^1/01355^; UAS-bru1-GFP* egg chamber, stained for Cyc B via IF. Images are deconvolved, XY max-intensity Z-projections 4 optical slices (0.3µm each). Scale bar, 10μm.

To determine whether this rescue is related to changes in aberrant Cyclin expression, we assessed the ectopic expression of CycB. Although ectopic CycB protein is still detected with UAS-Bru1-GFP overexpression, its levels are reduced (Figure 4C). It is of note that the UAS-Gal4 system is known for generating mosaic transgene expression. This results in variable levels of Bru1-GFP across nurse cells. Strikingly, we observed an inverse relationship between Bru1-GFP and CycB expression: nurse cells with higher Bru1-GFP levels exhibit lower CycB expression, while those with lower Bru1-GFP levels show increased CycB expression (Figure 4C, D). This suggests that Bru1 plays a role in the translational repression of *cycB* mRNA, despite the lack of detectable direct binding via UV crosslinking [7].

### Bru1 and Cup mediate accumulation of *cycA* and *cycB* mRNAs with Me31B

We found that *cycA* and *cycB* mRNAs associate with Bru1 and Cup within large cytoplasmic condensates, and such interactions are critical for the transcripts’ stability and translational repression. Both Bru1 and Cup are known components of P-bodies, membraneless cytoplasmic organelles, where mRNAs are stored in a translationally repressed state. Me31B, another core P-body component, forms a complex with Cup and is implicated in mRNA translational repression. A global analysis of early embryos revealed that Me31B associates with the majority of maternal mRNAs [35–38].

This led us to further explore the link between *cycA* and *cycB* mRNAs and Me31B. Using STED microscopy, we assessed the colocalization of these transcripts in egg chambers co-expressing Me31B-GFP and Cup-YFP, and found that larger mRNA puncta accumulated within Me31B/Cup-bodies, while single mRNA copies were not associated with either protein (Figure 5A), mimicking the results with Cup-YFP alone (Figure 2E, F). To confirm their localization in P-bodies, we performed combined smFISH-IF experiments, additionally detecting Bru1 with *cycA*/*cycB* mRNPs in sectioned tissue. As anticipated, all three proteins (Me31B, Cup, and Bru1) and both mRNAs colocalized in large cytoplasmic complexes, suggesting that *cycA* and *cycB* mRNAs are indeed recruited and stored in P-bodies (Figure 5B, S5C).

**Figure 5.**
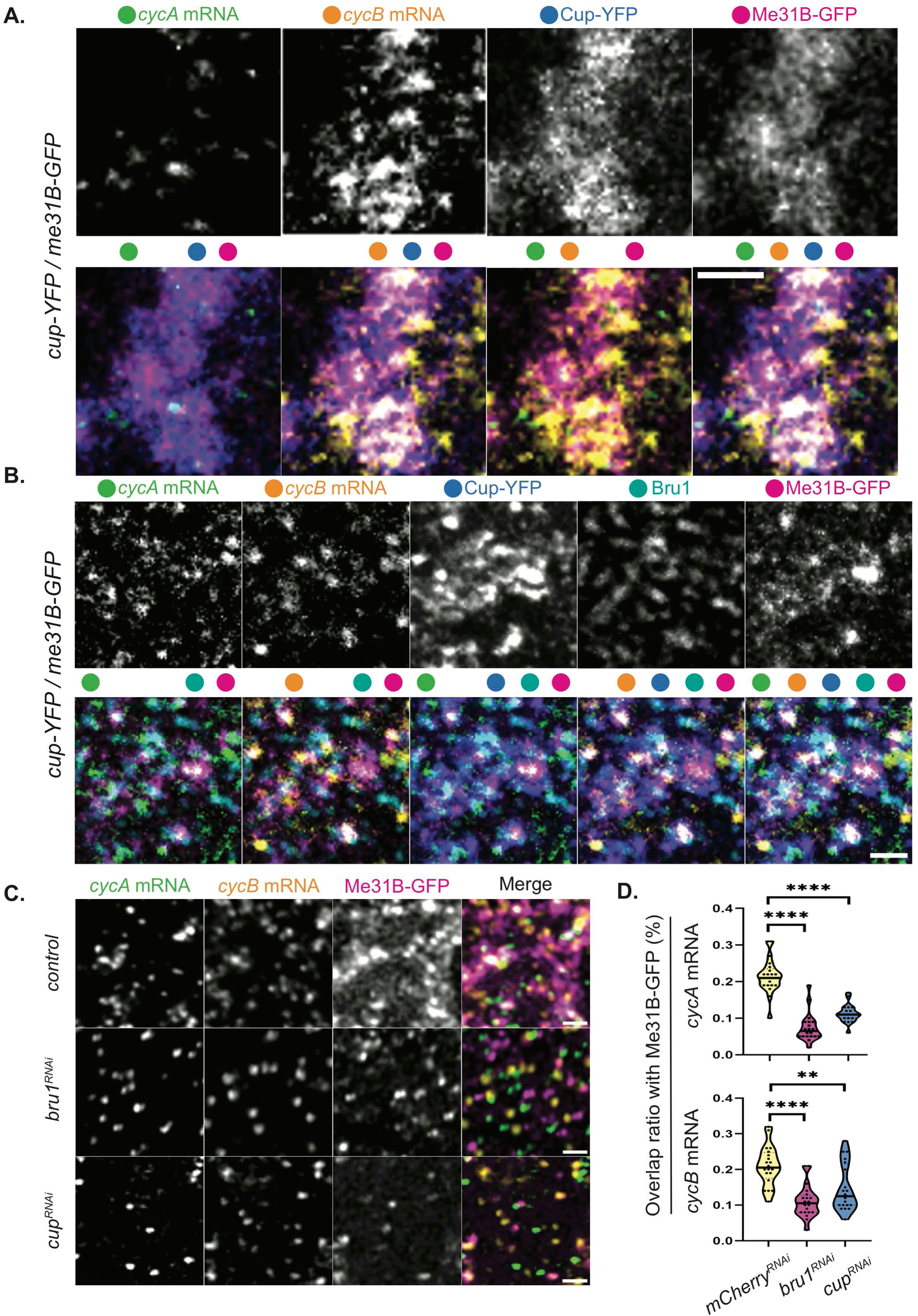
*CycA* and *cycB* mRNAs accumulate in Me31B-bodies in a Bru1 and Cup-dependent manner. **(A)** Co-visualization of *cycA* and *cycB* mRNAs with smFISH via STED in the nurse cells of a sectioned mid-oogenesis egg chamber expressing Cup-YFP and Me31B-GFP. Colored dots indicate individual channels merged per image. Images are deconvolved, XY max-intensity Z-projections of 3 optical slices (0.1µm each). Scale bar 1μm. **(B)** Co-visualization of *cycA* and *cycB* mRNAs via smFISH and Bru1 protein via IF in the nurse cells of a sectioned egg chamber at mid-oogenesis, expressing Cup-YFP and Me31B-GFP. Colored dots indicate the channels merged in the images. Images are deconvolved, XY max-intensity Z-projections of 5 optical slices (0.2µm each). Scale bar 1μm. **(C)** Co-visualization of *cycA* and *cycB* mRNAs via smFISH in stage 5 egg chambers expressing Me31B-GFP in indicated RNAi backgrounds. Images are deconvolved, XY max-intensity Z-projections of 4 (*control*), 5 (*cup^RNAi^*) and 5 (*bru1^RNAi^*) optical slices (0.3µm each). Scale bars, 1μm. **(D)** Overlap volume ratio analysis of *cycA* and *cycB* mRNAs with Me31B-GFP was performed in *control, cup^RNAi^* and *bru1^RNAi^*stage 2-4 egg chambers (****p<0.0001, **p=0.0035; *control* n=21, *cup^RNAi^*n=20, *bru1^RNAi^* n=20).

This finding prompted us to investigate why *cycA* and *cycB* mRNAs, despite their association with Me31B, undergo premature protein expression and mRNA degradation in the absence of Bru1 or Cup. We presumed that Me31B alone would be insufficient to maintain translational repression. To address this, we knocked down Bru1 or Cup in egg chambers expressing Me31B-GFP and evaluated the colocalization of *cycA* and *cycB* mRNAs with Me31B-GFP (Figure 5C). Notably, colocalization was significantly reduced; the overlap of *cycA* mRNA particles with Me31B particles decreased by 69% in *bru1^RNAi^* and 48% in *cup^RNAi^* egg chambers, while that of *cycB* mRNA was reduced by 48% and 38% in *bru1^RNAi^* and *cup^RNAi^*egg chambers, respectively (Figure 5D). These results indicate that while *cycA* and *cycB* mRNAs accumulate in Me31B-bodies, their localization depends on both Bru1 and Cup.

### Differential translational regulation of *cycA* and *cycB* mRNAs by Me31B

Given the severe reduction in *cycA* and *cycB* mRNA association with Me31B in Bru1– and Cup-depleted egg chambers, we questioned whether this diminished association drives the ectopic expression of CycA and CycB proteins. To address this, we knocked down Me31B in egg chambers expressing Me31B-GFP and Cup-YFP and assessed CycA and CycB expression. Using the same Gal4 driver as in the Bru1 and Cup knockdowns, we observed that these egg chambers progressed to stage 10/11 without mitotic spindle formation, exhibited a disrupted actin cytoskeleton and contained multiple nurse cell nuclei within a shared cytoplasm (Figure 6A, S6A). Me31B knockdown caused *cycB* mRNA derepression during early oogenesis, lasting until stage 6, though this derepression was less pronounced than in *cup^RNAi^*egg chambers (Figure 6A). Notably, no ectopic CycB protein could be detected beyond this stage in the *me31B^RNAi^* background. Cup and Bru1 levels were slightly reduced, and their cytoplasmic localization is altered, accumulating in distinct subcellular regions without forming the characteristic P-body condensates (Figure S6B, 3D-surface plots). Due to limited antibody penetration, Bru1 could not be fully visualized at stage 10 without sectioning the egg chambers, but some redistribution was detectable at stage 7 (Figure 6A, red arrow). Remarkably, no ectopic CycA protein expression was observed following Me31B knockdown (Figure 6B). These findings suggest that Me31B is not essential for maintaining the translational repression of *cycA* mRNA, but plays a role in *cycB* mRNA repression, potentially through the regulation of Cup and Bru1 expression levels and their cytoplasmic localization.

**Figure 6.**
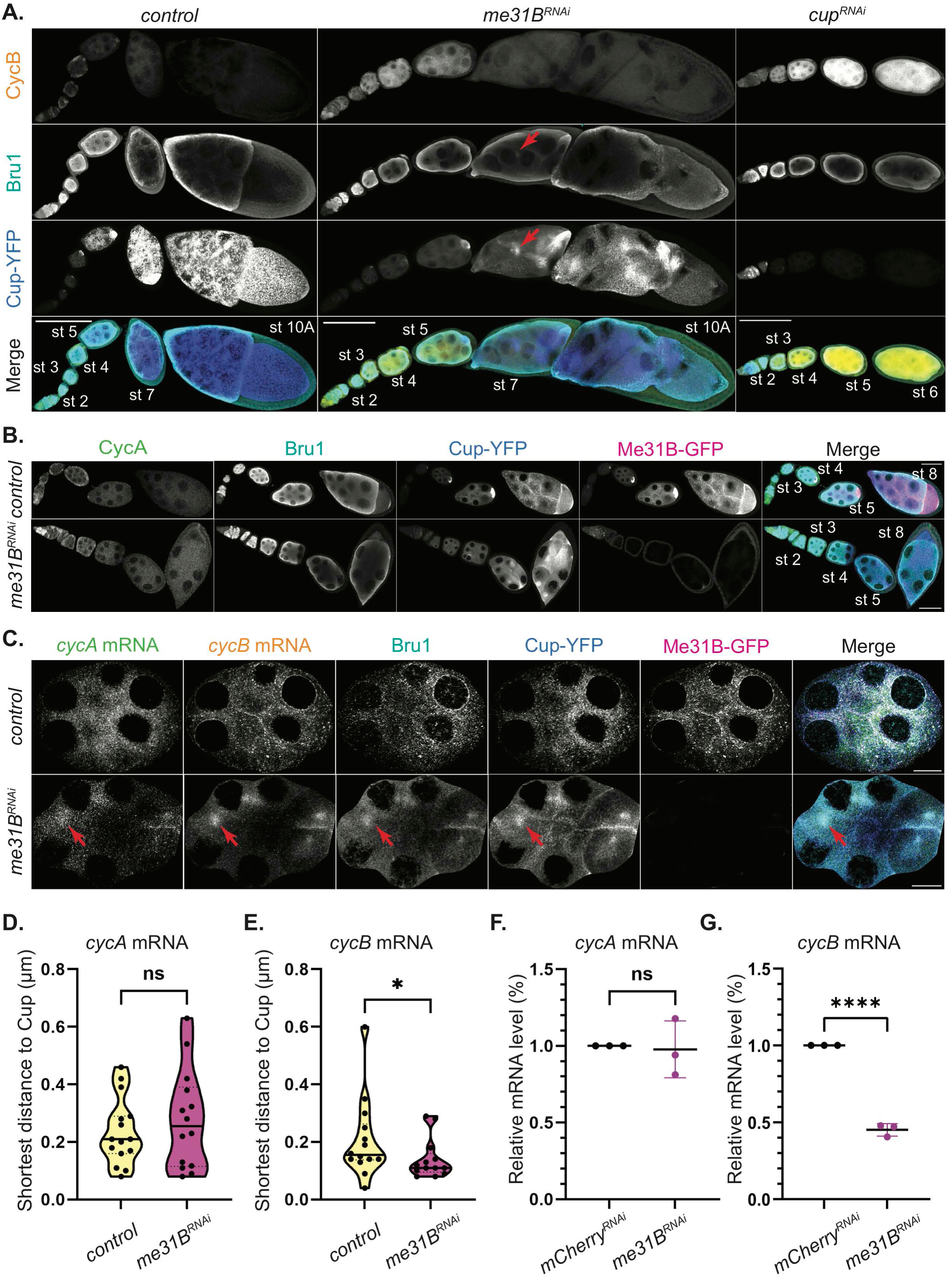
Me31B differentially regulates the translation of *cycA* and *cycB* mRNAs, but their association with Cup-YFP is independent of Me31B. **(A)** CycB and Bru1 visualized via IF in chains of egg chambers expressing Cup-YFP in the indicated RNAi backgrounds. Bru1 and Cup accumulation (red arrows). Images are deconvolved, XY max-intensity Z-projections of 15 (*control*), 13 (*me31B^RNAi^*) and 16 (*cup^RNAi^*) optical slices (1µm each). Scale bars, 100μm. **(B)** CycA and Bru1 visualized via IF in chains of egg chambers expressing Cup-YFP and Me31B-GFP in the indicated RNAi backgrounds. Images are deconvolved, XY max-intensity Z-projections of 6 (*control*) and 6 (*me31B^RNAi^*) optical slices (1µm each). Scale bars, 50μm. **(C)** Co-visualization of *cycA* and *cycB* mRNAs and Bru1 protein via smFISH and IF in sectioned mid-oogenesis egg chamber expressing Cup-YFP and Me31B-GFP in *control* and *me31B^RNAi^* backgrounds. *cycA* mRNA, *cycB* mRNA, Bru1 and Cup accumulation (red arrows). Images are deconvolved, XY max-intensity Z-projections of 5 (*control*) and 9 (*me31B^RNAi^*) optical slices (0.2µm each). Scale bars, 10μm. **(D)** Shortest distance analysis of *cycA* mRNA to Cup-YFP was performed in *control* and *me31B^RNAi^* backgrounds in stages 6-8 egg chambers (NS; *control* n=15, *me31B^RNAi^* n=14). **(E)** Shortest distance analysis of *cycB* mRNA to Cup-YFP was performed in *control* and *me31B^RNAi^* backgrounds in stages 6-8 egg chambers (*p=0.049; *control* n=14, *me31B^RNAi^* n=13). **(F)** RT-qPCR quantification of endogenous *cycA* mRNA normalized to *rp49* mRNA (mean ± SEM; NS; *mCherry^RNAi^* n=3, *me31B^RNAi^* n=3). **(G)** RT-qPCR quantification of endogenous *cycB* mRNA normalized to *rp49* mRNA (mean ± SEM; ****p<0.0001; *mCherry^RNAi^* n=3, *me31B^RNAi^* n=3).

### *cycA* and *cycB* mRNAs association with Cup-YFP is independent of Me31B

Given the absence of detectable CycA and only limited ectopic CycB protein in *me31B^RNAi^* egg chambers, we hypothesized that the mRNP complex retained its integrity, thereby maintaining the translational repression of both transcripts. To test this, we analyzed the colocalization of *cycA* and *cycB* mRNAs with Cup and Bru1 in *me31B^RNAi^* egg chambers. We conducted smFISH-IF experiments on sectioned egg chambers to overcome the limitations posed by antibody penetration. Our results revealed that Bru1, Cup and *cycA* and *cycB* mRNAs exhibit altered cytoplasmic distribution, no longer forming large cytoplasmic condensates, yet they accumulate together in subregions within the cytoplasm (Figure 6C, red arrows, and S6B, 3D-Surface plots). In some cases, these accumulations were observed to occur near the ring canals, likely due to impaired mRNA transport between nurse cells that results from the breakdown of the actin cytoskeleton (Figure S6C, red arrows). Further analysis of the egg chambers beyond stage 7, showed that the association between *cycA* mRNA and Cup was not significantly altered (Figure 6D). Interestingly, the observed distance between *cycB* mRNA particles and Cup particles decreased, indicating that their association was not only preserved but became more tightly coupled (Figure 6E). To determine whether Me31B plays a role in maintaining wild-type mRNA levels of *cycA* and *cycB*, we performed RT-qPCR. While *cycA* mRNA levels remained unchanged (Figure 6F), *cycB* mRNA levels showed a significant reduction (55%) (Figure 6G).These findings suggest that although *cycA* mRNA accumulates with Me31B, this interaction is not essential for its post-transcriptional regulation. In contrast, *cycB* mRNA stability appears to depend on P-bodies for protection from degradation, though P-bodies likely serve to fine-tune its translational repression, which is primarily mediated by Bru1 and Cup.

## Discussion

The precise regulation of the cell cycle relies on the proper expression and degradation of Cyclins, including CycA and CycB, two proteins essential for the initiation and progression of mitosis (reviewed in [2]). Aberrant expression of either protein can result in developmental arrest and cancer, as tumor-associated mutations often target CDK/Cyclin complexes, leading to uncontrolled cell division. Thus, it is critical to deepen our understanding of how these mRNAs are regulated.

We found that the knockdown of either Bru1 or Cup in the *Drosophila* germline led to the ectopic expression of both CycA and CycB proteins during oogenesis. However, while the CycB protein was detected immediately following the knockdown of either Bru1 or Cup, robust CycA expression was only observed beginning at stage 4, which coincides with the re-entry into mitosis in the knockdown egg chambers. This suggests that both proteins must reach a threshold to initiate mitotic division, or alternatively, that CycA is necessary for mitosis while CycB alone is insufficient. The latter model is more plausible, as previous studies have shown that CycA-deficient embryos that accumulate CycB still fail to enter mitosis [39]. Another possible explanation for the onset of mitosis at stage 4 is that nurse cells, at this stage, exit the endocycle, and their chromosomes undergo partial mitotic chromatin condensation, possibly rendering the nurse cells more susceptible to mitotic re-entry [5]. These explanations are not mutually exclusive and may likely contribute together to the observed phenomenon. Our findings reveal that, while both CycA and CycB proteins are ectopically expressed in the absence of Bru1 and Cup, their distinct expression patterns suggest differential regulatory mechanisms governing their mRNAs.

For *cycB* mRNA, the alternative splicing of its 3’ UTR is a likely regulatory mechanism, resulting in the expression of a short and long isoform. The short transcript is expressed during early oogenesis (< stages 6/7), when Bru1 levels are high and *cycB* mRNA is translationally repressed, while the long transcript is expressed during late oogenesis (> stages 9), when Bru1 is no longer detectable [4, 7]. Our analysis, conducted prior to the expression of the long transcript, confirmed that Bru1 colocalizes with the short isoform of *cycB* mRNA. Interestingly, despite the absence of canonical BREs in the *cycB* transcript we still observed a clear colocalization between Bru1 and *cycB* mRNA. This suggests that Bru1 may recognize and bind *cycB* mRNA through non-canonical sites or an indirect mechanism, highlighting a potentially broader role for Bru1 in *cycB* mRNA regulation.

Two models have been proposed to explain *cycB* mRNA translational derepression in the *bru1* mutant. The first posits that CycA expression triggers the CycA/CDK1 complex, which subsequently relieves the inhibition of CycB, making CycB protein expression a secondary effect of CycA expression rather than being directly regulated by Bru1. However, we find this model unlikely, as the CycB protein consistently appears in earlier stages of oogenesis than CycA. We favor the second model, which suggests that Bru1 regulates *cycB* mRNA in a BRE-independent manner, similar to its regulation of *germ cell-less* (*gcl*) mRNA [40]. Whether this interaction is direct or mediated by another protein, potentially Cup, remains unclear.

Further investigation is required to determine whether the alternative splicing of the 3’ UTR affects Bru1/Cup binding by altering mRNA folding or if the longer UTR recruits a protein that inhibits Bru1/Cup-mediated repression or eliminates their binding. Additionally, since Bru1 has been implicated in splicing regulation in muscle tissue, it would be intriguing to investigate whether Bru1 also regulates *cycB* mRNA splicing. Moreover, it would be valuable to explore whether the splicing of other Bru1 target mRNAs is altered in *cup* knockdown egg chambers, where Bru1 accumulates in large condensates around the nurse cell nuclei [41, 42].

A key piece of evidence supporting Bru1’s role in *cycB* mRNA translational regulation is the suppression of CycB protein expression in *cup* mutants when Bru1 is overexpressed. While the underlying mechanism remains unclear, it is possible that in *cup* mutant egg chambers, where Cup levels are significantly reduced but not entirely absent, Bru1 overexpression increases the formation of silencing particles, thereby rescuing egg chamber development. Alternatively, in the complete absence of Cup, elevated Bru1 levels may promote Bru1 dimerization, resulting in the formation of silencing complexes, similar to those observed in *oskar* mRNA translation repression [43].

How Bru1 is recruited to *cycB* mRNA remains an unresolved question, and we propose several potential mechanisms for this interaction. First, Cup, which is known to directly bind mRNAs, could act as a mediator, facilitating the recruitment of Bru1 to *cycB* mRNA [44]. Second, Bru1 may bind *cycB* mRNA in a BRE-independent manner, similar to its interaction with *gcl* mRNA [40]. Third, Bru1 could be recruited to *cycB* mRNA through interaction with another protein, and in *cup* mutants —where Bru1 levels are reduced— only the overexpression of Bru1 may provide sufficient levels to restore *cycB* mRNA repression. While these models suggest plausible mechanisms for *cycB* mRNA regulation, the precise pathway remains to be clarified through future studies.

In the embryo, *cycB* mRNA is translationally repressed through the binding of Pumilio and Nanos to the 3’ UTR, with Nanos being the essential factor. Nanos recruits the CCR4-NOT complex specifically in germ cells, where *cycB* mRNA is deadenylated, but not degraded. Additionally, Nanos directly interacts with Cup, colocalizing in both the germarium and the primordial germ cells of the embryo [45, 46]. Nanos levels are high in the germarium, but rapidly decrease as the egg chamber exits this region [47]. Interestingly, both Nanos and Cup exert their translational repression by recruiting the CCR4-NOT complex and are involved in the sequential regulation of another mRNA, *pgc* mRNA, during oogenesis. Before differentiation in the germarium, Pumilio and Nanos repress *pgc* mRNA, while after differentiation, this role is taken over by Bru1 and Cup [28]. Whether *cycB* mRNA undergoes similar regulation remains to be determined, yet we believe this mechanism is highly probable. Supporting this hypothesis, in the embryo, *cycB* mRNA regulation requires not only Nanos, but also an Oskar-dependent factor, which we predict could be Cup [45].

The formation of the translationally repressive *cycB* mRNP is essential not only for *Drosophila* development but also for *Xenopus*, zebrafish and mouse oocyte development, underscoring its evolutionary conservation [48, 49]. The canonical model for this regulation involves a 3’ UTR-binding protein recruiting an eIF4E-binding protein, which in turn inhibits the formation of the translational initiation complex. This mechanism has been demonstrated for numerous mRNAs, particularly in germ cell mRNPs (reviewed in [50, 51]). Additionally, *cycB*1 mRNA translational control is regulated through the assembly and disassembly of mRNA granules in zebrafish and mouse oocytes [48]. These mRNPs can subsequently form higher-order complexes via LLPS, contributing to the formation of P-bodies [15, 52, 53].

P-bodies, where mRNPs accumulate, are cytoplasmic sites for mRNA storage and translational repression [15, 19, 52]. Both Bru1 and Cup are known to localize within P-bodies, but whether their accumulation into P-bodies is essential for maintaining mRNAs in a translationally repressed state remains controversial [12, 27, 53]. Recent research has shown that cyclic mRNAs dynamically accumulate in P-bodies during the cell cycle, with mRNAs in P-bodies being uncoupled from their cytoplasmic expression, suggesting a cell cycle-dependent targeting of mRNAs to P-bodies [54].

Through our study, we show that *cycA* and *cycB* mRNAs are both regulated by Bru1 and Cup and both accumulate in P-bodies, but exhibit differential regulation. For *cycA* mRNA, maintaining the integrity of the primary mRNP complex is essential for its post-transcriptional regulation, independent of P-body association. In contrast, *cycB* regulation is more complex: translational repression is not fully sustained, and *cycB* mRNA levels are significantly reduced in Me31B knockdowns. These findings suggest that *cycB* mRNA recruitment to P-bodies is important for its regulation. However, since the changes in expression were not as pronounced as in the Bru1 or Cup knockdown egg chambers, P-body recruitment likely serves to fine-tune *cycB* mRNA translational regulation, while maintaining the integrity of the primary mRNP complex is key. The exact mechanism behind this differential regulation remains unknown, but identifying the variations in mRNP composition will likely provide crucial insights. Finally, given our recent discovery that ER exit sites contribute to P-body organization in the case of Me31B, but not for Cup, it would be intriguing to explore the role of ER exit sites in mediating the association of *cyclin* mRNAs with Me31B or Cup [55].

We highlight that the formation of the mRNP complex is critical for maintaining the repression and stability of both *cycA* and *cycB* mRNAs. While P-body accumulation is not required for *cycA* regulation, it is important for proper *cycB* regulation. This finding underscores the complexity of P-body involvement in mRNA regulation and suggests there is much more to learn about their roles. Future research should aim to uncover the mechanisms behind this differential regulation, which could provide broader insights into P-body functions or P-body subclasses. Our results show that P-bodies modulate mRNA regulation in a transcript-specific manner rather than through a uniform mechanism.

## Methods

### Fly husbandry

*Drosophila melanogaster* stocks were maintained on standard cornmeal agar food at 25°C. Prior to dissection, female flies were fed yeast paste in grape vials for 2-5 days. Fly stocks were obtained from Bloomington *Drosophila* Stock Center. Gal4-inducible TRiP (RNAi) lines: *mCherry* (BL #35785), *cup* (BL #35406), *bru1* (BL #35394 and BL #54812), *me31B* (BL #38923 and BL#33675). Mutant lines: *cup^01355^* (BL #12218) and *cup^1^* (BL #4978). Maternal Gal4 driver line *matalpha4-GAL-VP16 (V37)*(BL#7063) was used throughout the study. Fluorescently tagged lines: *bru1-GFP* (BL #60144) and *me31B-GFP* (BL #51530). We acquired *cup-YFP* (DGRC 115-161) from the Kyoto *Drosophila* Stock Center. *UAS-bru1-GFP* fly line – gift from Dr. P.M. Macdonald (The University of Texas at Austin).

### Sectioning of egg chambers

Ovaries were dissected into 2% PFA in PBS and fixed for 10 min. They were washed 3 x 10 min in PBST and 1 x 5 min in 0.1M glycine pH 3, and stored in 30% sucrose overnight at 4°C. The following day, they were flash frozen in O.C.T Compound (Tissue-Tek) before 25µm sections were prepared on a cryotome.

### Immunofluorescence

Ovaries were dissected and fixed in 2% PFA in PBS for 10 min. Egg chambers were washed 3 x 5 min in PBST (0.3% Triton X-100), permeabilized and blocked for 2 hrs in PBS with 1% Triton X-100 and 1% BSA, and incubated with primary antibodies overnight at room temperature with rocking.

Followed by 3 x 15 min washes in PBST and incubation with fluorescently labeled secondary antibodies (1:1000; DyLight 550 and DyLight 650; ThermoFisher Scientific). Followed by 3 x 15 min washes in PBST and mounting in ProLong Diamond Antifade Mountant (Life Technologies) or in a mixture of 75:25 RapiClear 1.47 (SUNJin Lab):Aqua-Poly/Mount (Polysciences) for clearing the egg chambers. Antibodies used: mouse anti-CycA A12 – deposited to the DSHB by Lehner, C.F. (DSHB Hybridoma Product A12)(1:100); mouse anti-CycB F2F4 –deposited to the DSHB by O’Farrell, P.H. (DSHB Hybridoma Product F2F4) (1:100); mouse anti-Cup (1:1000) and mouse anti-Me31B (1:1000) – gifts from Dr. A. Nakamura (Institute of Molecular Embryology and Genetics, Kumamoto University); rabbit-anti-Bruno 1 (1:5,000) – gift from Dr. M. Lilly (National Institutes of Child Health and Development, NIH); rabbit-anti-Bruno 1 (1:4,000) – gift from Dr. P.M. Macdonald (The University of Texas at Austin); F-actin stain: Phalloidin Alexa Fluor 647 (1:500; Life Technologies), nuclear membrane stain: wheat germ agglutinin CF^Ⓡ^405S and CF^Ⓡ^555 (1:400; Biotium). Actin and nuclear membrane stains were added together with the secondary antibody incubation.

### Tubulin staining

Ovaries were dissected into BRB80 (0.5M K-PIPES pH 6.8, 2M MgCl_2_, and 0.5M K-EGTA) and permeabilized in 1% Triton X-100 in BRB80 without rocking. They were rinsed in BRB80 and fixed with 2% PFA for 10 min. Egg chambers were washed 3 x 10 min in PBST followed by 1 x 10 min in 2X SSC 10% formamide, and incubated overnight in mouse anti-–Tubulin AA4.3-s deposited to the DSHB by Walsh, C. and anti-–Tubulin 12G10 deposited to the DSHB by Frankel, J. / Nelsen, E.M (1:100; Developmental Studies Hybridoma Bank). Egg chambers were washed 3 x 10 min followed by 2 hr incubation in a fluorescently labeled secondary antibody (1:1000; DyLight 550 or DyLight 650; ThermoFisher). Egg chambers were washed 3 x 20 min in PBST and were fixed again in 2% PFA, washed 3 x 5 min in PBST and mounted, as described above.

### Single-molecule RNA FISH (smFISH) in ovaries

smFISH were performed as previously described [56] with the following modifications. Ovaries were dissected and fixed in 2% PFA in PBS for 10 min. Egg chambers were washed 3 x 5 min in PBST (0.3% Triton X-100). After fixation, egg chambers were pre-hybridized in wash buffer (2X SSC, 10% formamide) and incubated with the *cycA*-570 and *cycB*-610 probes (1:100) (Supplemental_Methods_S1) overnight at 37°C, followed by washing 3 x 10 min in wash buffer and mounting with ProLong^TM^ Diamond Antifade Mountant (Life Technologies) on a glass slide using a #1.5 coverslip glass.

### Single smFISH and combined smFISH-IF in sectioned egg chambers

Sectioned egg chambers were washed and re-hybridized in PBS 2 x 10 min and in 2X SSC for 10 min followed by pre-hybridization in wash buffer (2X SSC, 10% formamide) for 10 min. Sections were incubated with the *cycA*-570 and *cycB*-610 probes (Supplemental_Methods_S1) for 8 hours or overnight at 37°C, followed by washing with wash buffer and mounting with ProLong^TM^ Diamond Antifade Mountant (Life Technologies) on a glass slide using a #1.5 cover glass. Combined smFISH-IF was carried out following 8 hours of incubation with the smFISH probes. Sections were washed in the wash buffer for 10 min and then 0.05% Triton X-100 in 2X SSC for 10 min. Followed by incubation with primary antibodies overnight at room temperature in 0.05% Triton X-100, 0.2% BSA, 2X SSC. After 3 x 10 min washes with 0.05% Triton X-100 in 2X SSC, the egg chambers were incubated with fluorescently labeled secondary antibodies (1:1000; DyLight 550 or DyLight 650; ThermoFisher) for 2 hr at room temperature in 0.05% Triton X-100, 0.2% BSA, 2X SSC, and were washed 3 x 10 min with 0.05% Triton X-100 in 2X SSC and mounted with ProLong^TM^ Diamond Antifade Mountant (Life Technologies) on a glass slide using a #1.5 coverslip glass.

### Microscopy

All imaging was performed on a TCS SP8 Laser Scanning Microscope (Leica Microsystems) equipped with a white light laser (470-670nm), a solid-state laser (405nm), and a STED 660nm CW high intensity laser. For all laser scanning confocal imaging, the HyD detectors were used in photon counting mode, and 40X/1.4 and 63X/1.4 oil objectives were used. For all STED imaging, the 100X/1.4 oil objective was used, optical Z slices were 0.1µm, and zoom was set to 6X. STED samples were prepared from 25µm ovary sections. Optical sections were acquired using an automated XYZ-piezo stage. Acquisition software: Leica LAS-X. Images were merged using the smooth option with Leica’s ‘Image merger’.

### Imaging analysis

Identical image acquisition was used for the control, as well as each experimental slide. For all analyses, images were taken from 3 slides each prepared from an independent fly cross. All image files were saved as 16-bit data files after deconvolution with Leica’s ‘Lightning’ module. Figure images were processed using Fiji/ImageJ (NIH) [57, 58]. Quantification was performed using Imaris Microscopy Image Analysis software (Oxford Instruments). Imaris ‘Surface detection’ module was used to identify the objects for the volume overlap and shortest distance calculations. Thresholding and sensitivity were carefully determined, and the mean value of each image was used for the statistical calculations. Statistics were calculated using Mann-Whitney statistical tests on Prism 8 Software (GraphPad).

### Western blot analysis

Ten ovaries for each genotype were dissected directly into 95µl of 2X Laemmli Sample Buffer (Bio-Rad) supplemented with 5µl of BME and immediately mechanically lysed. They were heated at 95°C for 10 min and centrifuged for 10 min at 10,000 RCF at 25°C. Lysates were loaded onto a 10% acrylamide gel. Primary antibodies used: mouse anti-CycA (1:100), mouse anti-CycB (1:100), and rabbit anti-Tri-methyl-Histone (C42D8) (1:150,000; Cell Signaling Technology). Bands visualized using TrueBlot ULTRA secondary antibodies: anti-mouse and anti-rabbit Ig HRP (1:50,000; Rockland) with SuperSignal West Femto Maximum Sensitivity Substrate (ThermoFisher Scientific).

### RNA isolation and RT-qPCR

Whole ovaries were dissected into 4°C PBS. Ovaries were mechanically lysed in TRIzol (ThermoFisher) to extract total RNA. RNA was washed with ethanol and eluted in RNAse-free water. Reverse Transcriptase reactions using 2.5µg total RNA were performed using Superscript IV kit (Life Technologies). Primers were designed using DRSC FlyPrimerBank and made by Integrated DNA Technologies (Supplemental_Methods_S2). RT-qPCR was performed with a Roche Lightcycler 480 (Roche Molecular Systems, Inc.). Each reaction contained 1µl of cDNA, 4µl of 10µM primers, and 5µl of PowerTrack™ SYBR Green Master Mix (ThermoFisher Scientific). Statistical significance was determined via t-test.

## Figure Legends

**Figure S1. Bru1 and Cup knockdowns lead to re-entry of nurse cells into the cell cycle and to the ectopic expression of CycA and CycB proteins**.

**(A)** –tubulin visualized via IF in stage 6 egg chambers expressing Cup-YFP and Bru1-GFP in the indicated backgrounds. Membrane is stained with WGA. Images are deconvolved, XY max-intensity Z-projections of 5 optical slices (0.3µm each). Scale bars, 10μm.

**(B)** –tubulin visualized via IF in a chain of egg chambers expressing Cup-YFP and Bru1-GFP in *cup^RNAi^* background. Membrane is stained with WGA. Red arrows indicate the re-formation of the nuclear membrane. Images are deconvolved, XY max-intensity Z-projections of 6 optical slices (0.3µm each). Scale bars, 50μm.

**(C)** Fluorescence intensity line plots of CycB protein. The region of interest and the direction of the line plots are outlined in the image.

**(D)** Fluorescence intensity line plots of CycA protein.The region of interest and the direction of the line plots are outlined in the image.

**(E)** Immunoblot against CycA and CycB. H3 – the loading control using lysates prepared from egg chambers in the indicated backgrounds.

**Figure S2. Bru1 and Cup form large complexes with *cycA* and *cycB* mRNAs in the nurse cell cytoplasm**.

**(A)** Visualization of *cycA* and *cycB* mRNAs in a chain of egg chambers. F-actin stain to outline the egg chambers. Images are deconvolved, XY max-intensity Z-projections of 10 optical slices (1µm each). Scale bar 50μm.

**(B)** Co-visualization via STED of *cycA* and *cycB* mRNAs in the nurse cells of a sectioned egg chamber at mid-oogenesis expressing Bru1-GFP and Cup-YFP. White box indicates the location of the zoomed in images in Figure 2A. Images are deconvolved, XY max-intensity Z-projections of 3 optical slices (0.1µm each). Scale bar 5μm.

**(C)** Co-visualization via STED of *cycA* and *cycB* mRNAs in the nurse cells of a sectioned egg chamber at mid-oogenesis expressing Bru1-GFP. Colored dots indicate the individual channels merged per image. White box indicates the location of the zoomed in images. Images are deconvolved, XY max-intensity Z-projections of 3 optical slices (0.1µm each). Scale bar 5μm.

**(D)** Co-visualization via STED of *cycA* and *cycB* mRNAs in the nurse cells of a sectioned egg chamber at mid-oogenesis expressing Cup-YFP. Colored dots indicate the individual channels merged per image. White box indicates the location of the zoomed in images. Images are deconvolved, XY max-intensity Z-projections of 3 optical slices (0.1µm each). Scale bar 5μm.

**Figure S3. Cup is necessary for the formation of *cycA* and *cycB* mRNPs**.

**(A)** Co-visualization of *cycA* and *cycB* mRNAs via smFISH in stage 6 egg chambers expressing Bru1-GFP in indicated backgrounds. White box indicates the location of the zoomed in images. Images are deconvolved, XY max-intensity Z-projections of 5 optical slices (0.3µm each). Scale bars, 10μm.

**(B)** Overlap volume ratio analysis of *cycA* and *cycB* mRNAs with Bru1-GFP performed in *control* and *cup1^RNAi^* stage 5-7 egg chambers (****p<0.0001; *control* vs *cup1^RNAi^*: n=11 and n=27).

**Figure S4. Bru1 overexpression rescues mitotic re-entry of *cup* mutant egg chambers**.

**(A)** Several ovariole chains of the indicated backgrounds with –tubulin visualized via IF. Images are deconvolved, XY max-intensity Z-projections of 5 (*UAS-bru1-GFP*), 6 (*cup^1/01355^*) and 3 (*cup^1/01355^; UAS-bru1-GFP*) optical slices (0.3µm each). Scale bars, 100μm.

**(B)** Chain of *wildtype* egg chambers expressing Bru1-GFP with CycB visualized via IF. Membrane (WGA). Images are deconvolved, XY max-intensity Z-projections of 5 optical slices. Scale bar 50μm.

**Figure S5. *cycA* and *cycB* mRNAs accumulate in Me31B-bodies in a Bru1 and Cup-dependent manner**.

**(A)** Co-visualization of *cycA* and *cycB* mRNAs via STED in the nurse cells of a sectioned egg chamber at mid-oogenesis, expressing Cup-YFP and Me31B-GFP. White box indicates the location of the zoomed in images in Figure 5A. Images are deconvolved, XY max-intensity Z-projections of 3 optical slices (0.1µm each). Scale bar 1μm.

**(B)** Co-visualization of *cycA* and *cycB* mRNAs and Bru1 via smFISH and IF in the nurse cells of a sectioned egg chamber at mid-oogenesis, expressing Cup-YFP and Me31B-GFP. White box indicates the location of the zoomed in images in Figure 5B. Images are deconvolved, XY max-intensity Z-projections of 5 optical slices (0.2µm each). Scale bar 1μm.

**(C)** Co-visualization of *cycA* and *cycB* mRNAs in stage 5 egg chambers expressing Me31B-GFP in indicated RNAi backgrounds. White box indicates the location of the zoomed in images in Figure 5C. Images are deconvolved, XY max-intensity Z-projections of 4 (*control*), 5 (*cup^RNAi^*) and 5 (*bru1^RNAi^*) optical slices (0.3µm each). Scale bars, 10μm.

**Figure S6. Me31B differentially regulates the translation of *cycA* and *cycB* mRNAs, but their association with Cup-YFP is independent of Me31B**.

**(A)** Co-visulaization of F-actin and DAPI in stage 9 egg chambers in *control* and *me31B^RNAi^* backgrounds.

**(B)** Co-visualization of *cycA* and *cycB* mRNAs via smFISH and F-actin in sectioned mid-oogenesis egg chamber expressing Cup-YFP and Me31B-GFP. F-actin was stained to visualize the membrane and the ring canals. *cycA* mRNA, *cycB* mRNA, and Cup accumulation at the ring canals in *me31B^RNAi^* (red arrows). Images are deconvolved, single optical slices. Scale bars, 10μm.

**(C)** Co-visualization of *cycA* and *cycB* mRNAs and Bru1 protein via IF in sectioned mid-oogenesis egg chambers expressing Cup-YFP and Me31B-GFP in *control* and *me31B^RNAi^*backgrounds. *cycA* mRNA, *cycB* mRNA, Bru1 and Cup accumulation (red arrows). 3D Surface plots depict the signal intensities (0-255) using 5_ramps LUT. Images are deconvolved, XY max-intensity Z-projections of 5 (*control*) and 9 (*me31B^RNAi^*) optical slices (0.2µm each). Scale bars, 10μm.

## ACKNOWLEDGEMENTS

We thank Dr. A. Nakamura (Riken Center for Developmental Biology) and Dr. M. Lilly (NIH) for the kind gifts of antibodies, as well as Dr. P.M. Macdonald (The University of Texas at Austin) for the kind gift of an antibody and the *D. melanogaster* line. We thank the BDSC Indiana, VDCR Vienna and DGRC Kyoto for providing *D. melanogaster* lines, as well as the TRiP at Harvard Medical School (NIH/NIGMS RO1-GM084947) for the transgenic RNAi stocks. We thank undergraduate students, Zara Kumar and Daniela Villafuerte, for their assistance with fly husbandry and the maintenance of stocks. We thank the Bioimaging Facility at Hunter College for access to the Leica TCS SP8 and Imaris –-Image Analysis Software. We thank Dr. P. Feinstein (Hunter College) for allowing us to use the Roche LightCycler instrument. This work was supported by the National Institute of Health (1SC1GM135132) and the National Science Foundation instrumentation award (1919829) to D. P. B.

## DATA AVAILABILITY

This study includes no data deposited in external repositories.

## AUTHOR CONTRIBUTIONS

**Bayer, V. Livia**: Methodology; Investigation; Formal analysis; Writing-original draft; Writing-review and editing. **Milano, N. Samantha**: Investigation; Writing-review and editing. **Harpreet Kaur**: Investigation; Writing-review and editing. **Bratu, P. Diana**: Methodology; Supervision; Funding acquisition; Project administration; Writing-review and editing.

