## Supplementary figures and images for "Post-transcriptional regulation of *cyclin A* and *cyclin B* mRNAs is mediated by Bruno 1 and Cup, and further fine-tuned within P-bodies"

### Supplemental Figure S1

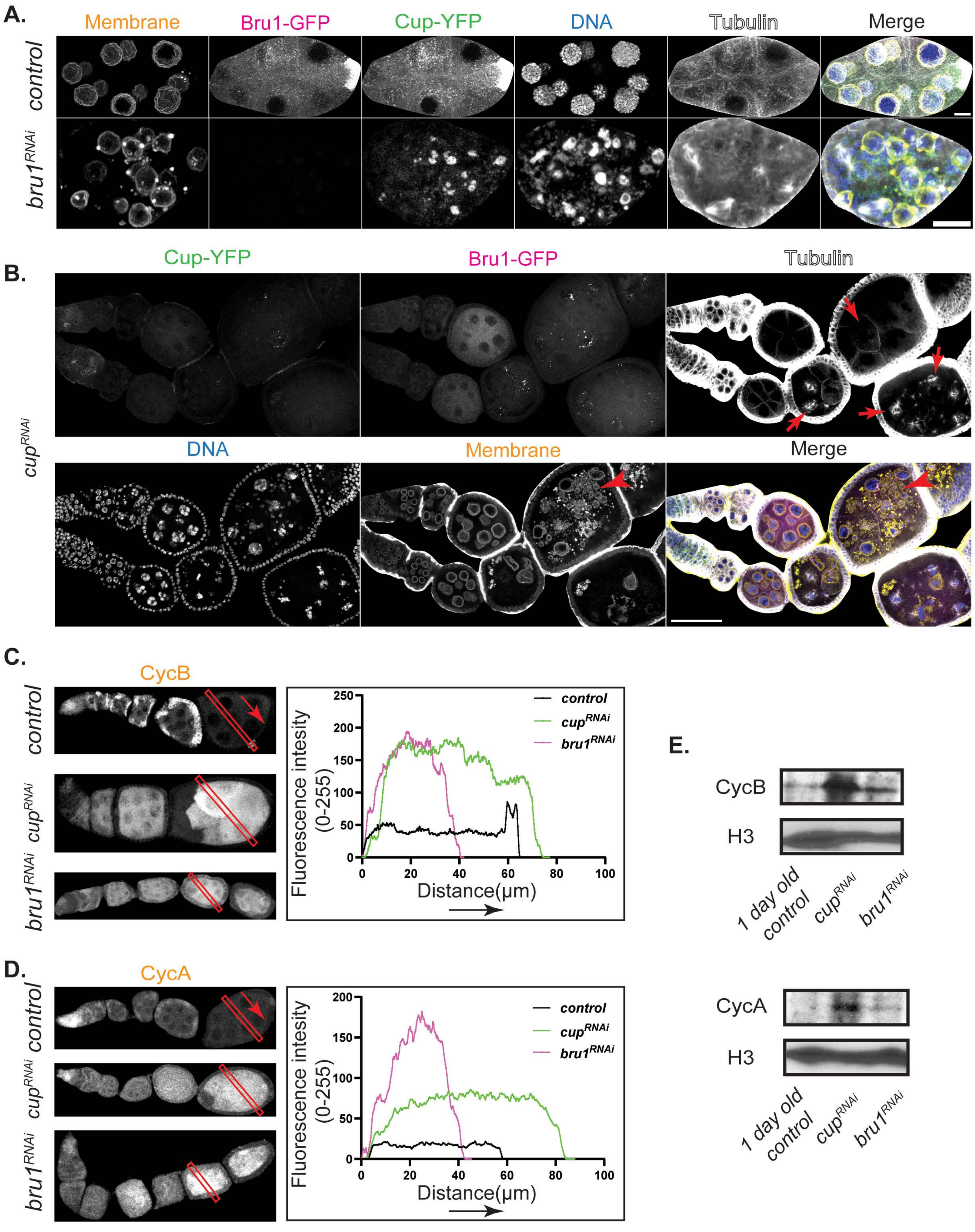

### Supplemental Figure S2

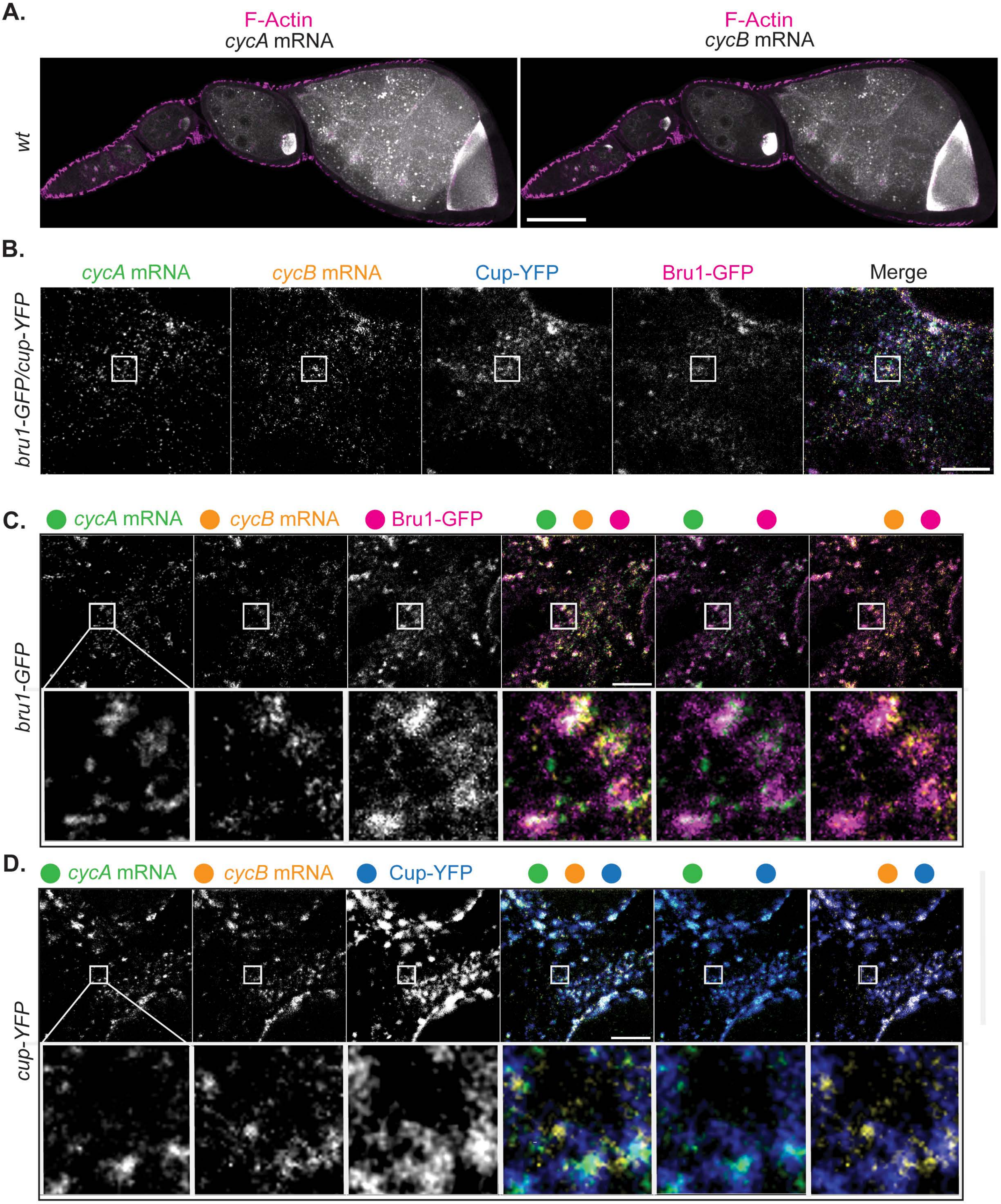

### Supplemental Figure S3

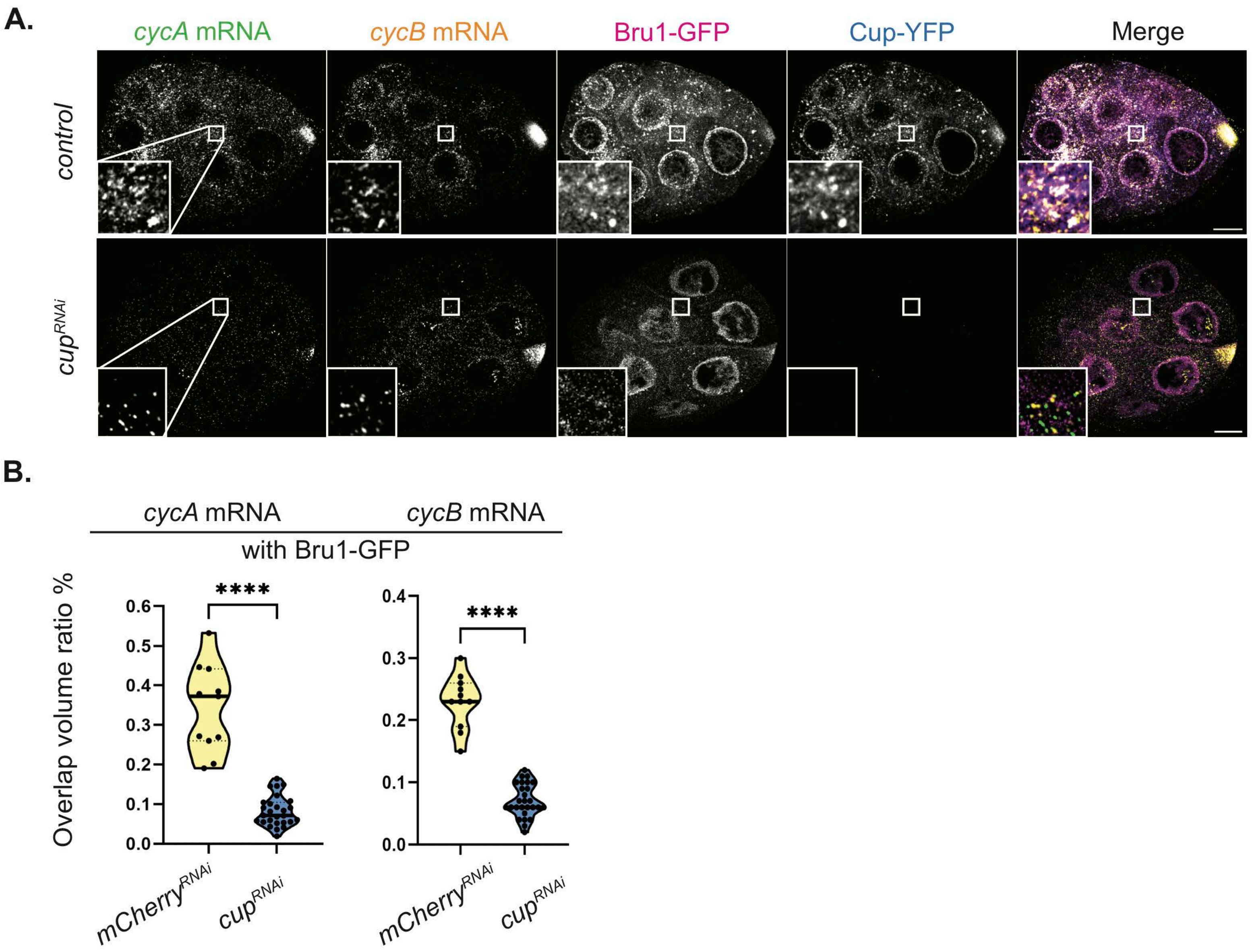

### Supplemental Figure S4

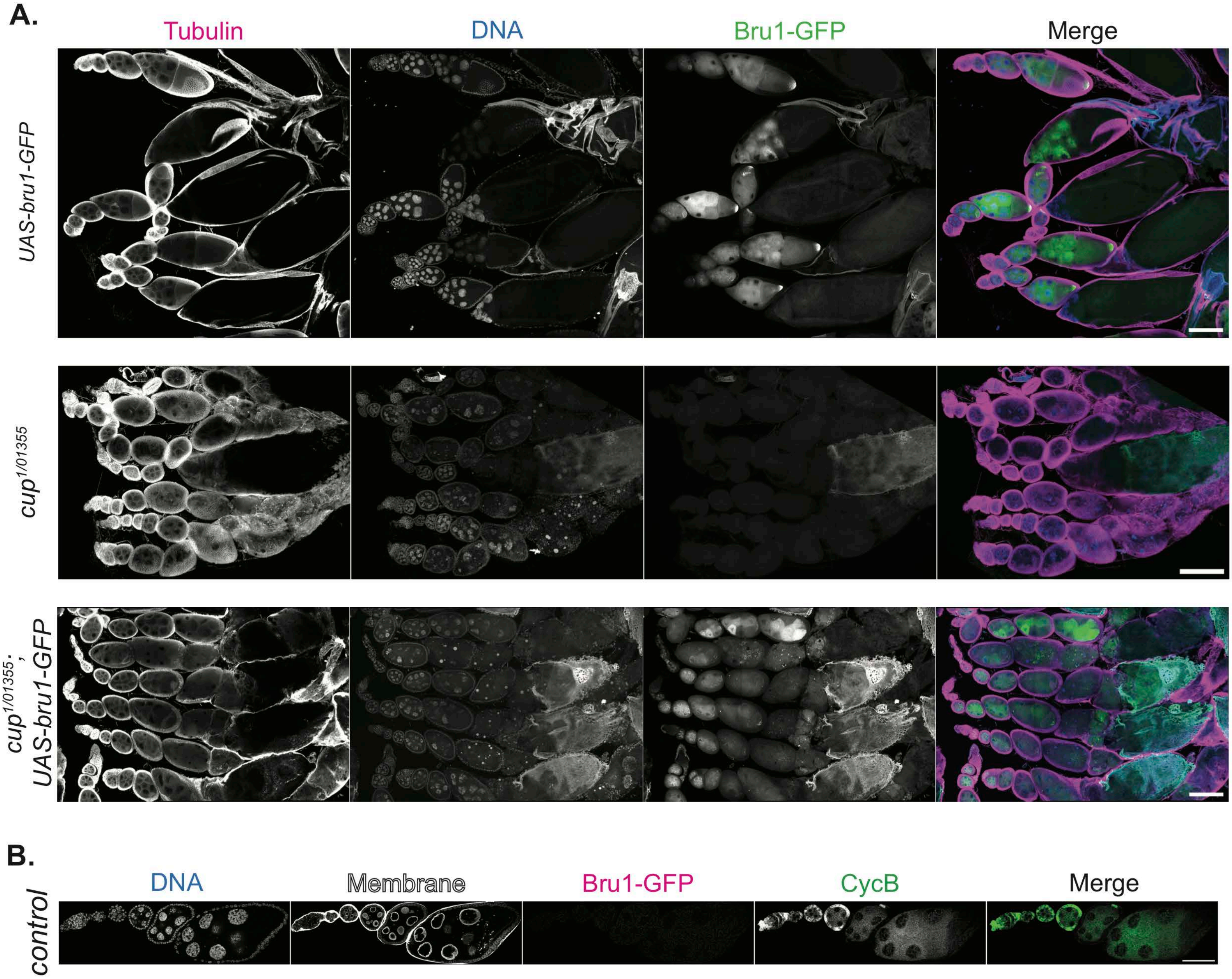

### Supplemental Figure S5

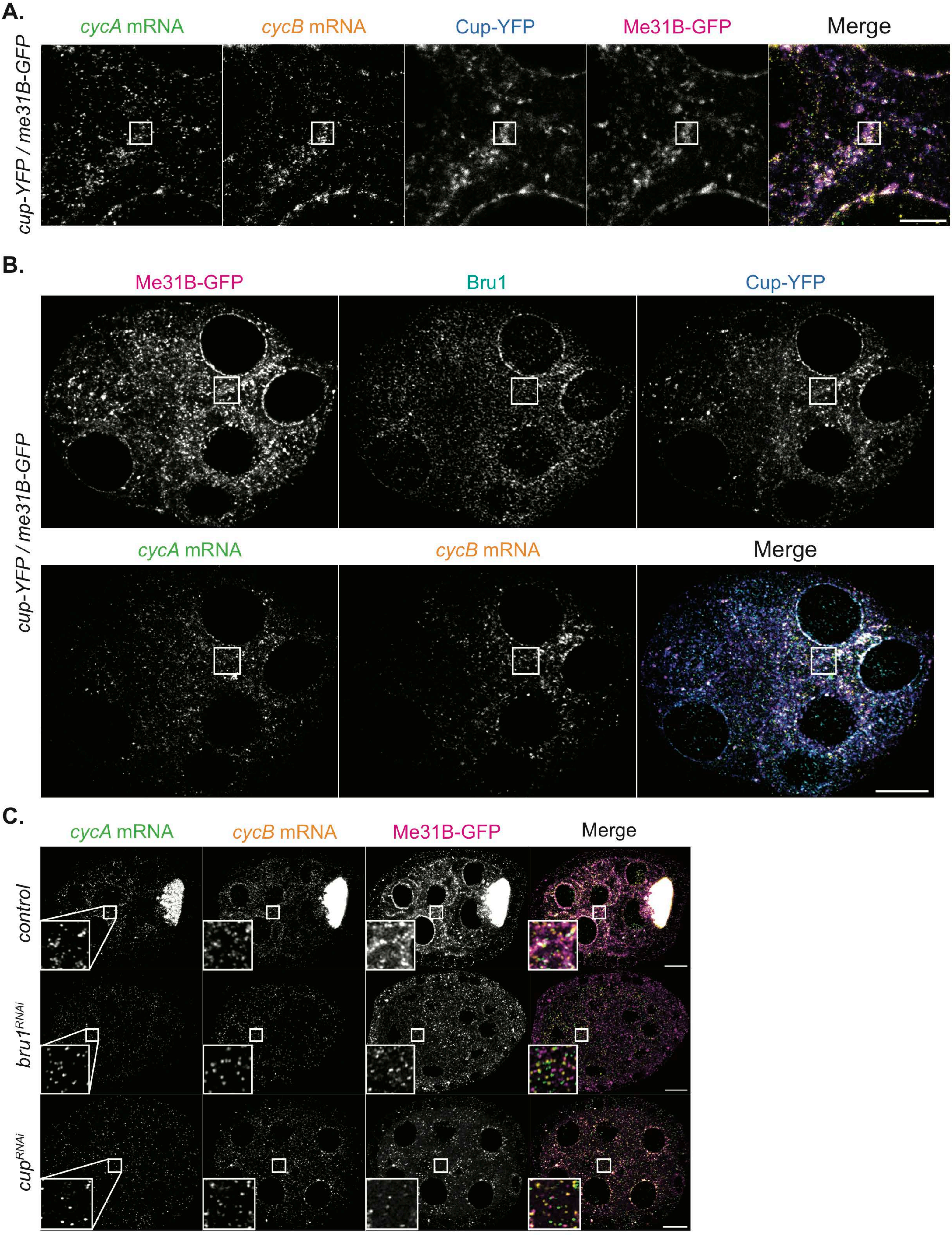

### Supplemental Figure S6

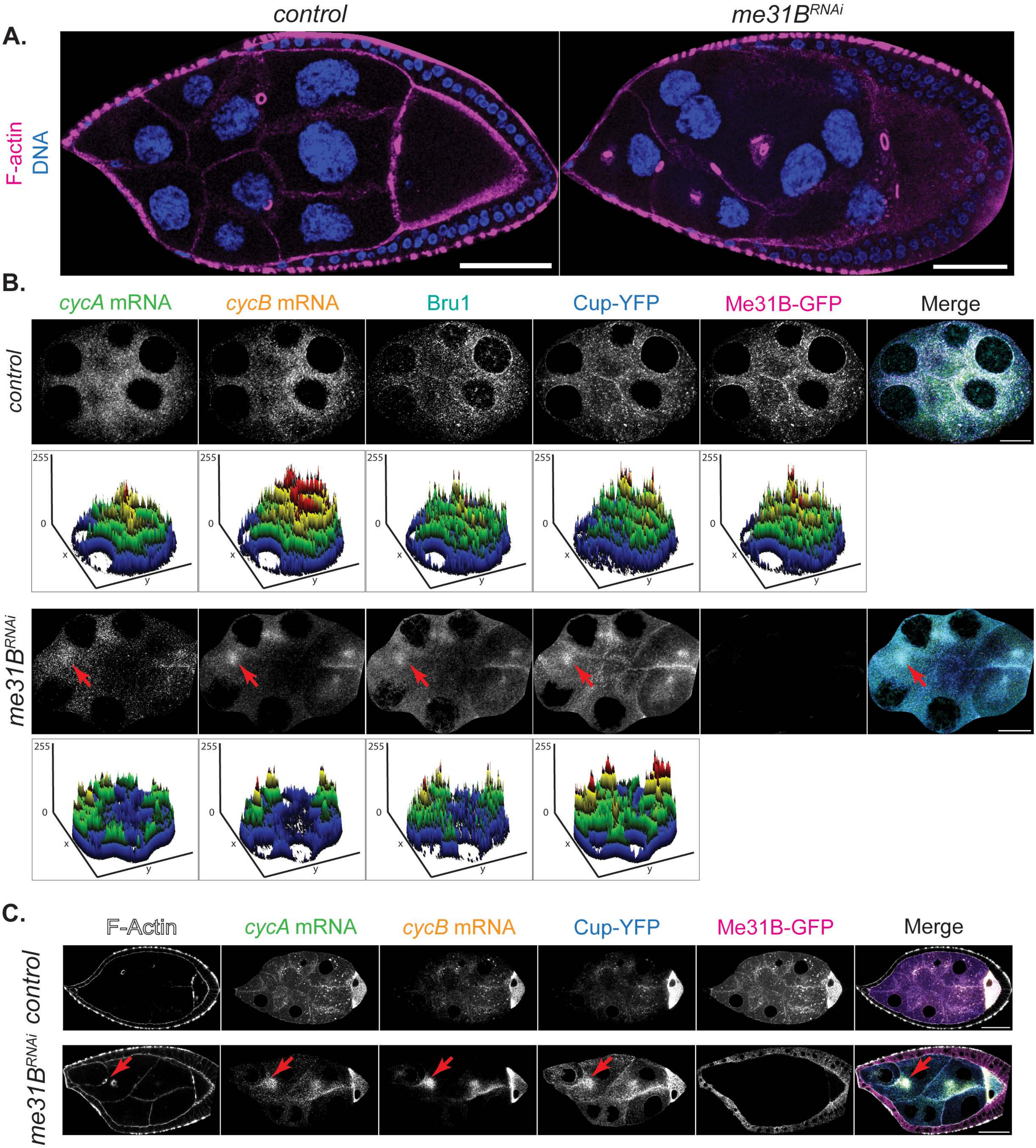
