## Supplemental Table S1 for "Post-transcriptional regulation of *cyclin A* and *cyclin B* mRNAs is mediated by Bruno 1 and Cup, and further fine-tuned within P-bodies"

**Table S1. Probe sequences used in smFISH experiments.**

| Probes | *cycB* mRNA | *cycA* mRNA |
|---|---|---|
| 1 | gattctgcaaatcgcccaag | gaatcttaatgcccggattc |
| 2 | ctgcgatgggacgacttatg | ctgcttggtgttcttcactc |
| 3 | ttgagatccttggagtcttt | attgagcacggcgaaattgg |
| 4 | cgaacgcaaaaacgccgaca | gacgcggcacattgttgttg |
| 5 | tcttggtcggaacgcgattg | gtcacggaagacaactttcc |
| 6 | actttagtgggttctacagt | actccacattctcatcaaca |
| 7 | ctcggaagaactggacttga | caccacattcgatttcttgg |
| 8 | tttaaggtgggctcgttcac | aaggtcttgaattgctccac |
| 9 | agcgacttcttcgacagatt | tatcgttgttgtcctcatac |
| 10 | caaagcggcacgcagtttgg | tctctttgtcgactagggag |
| 11 | aaactcccatcacgggtttg | agcaccgaatttcacgtcat |
| 12 | tactggttcccgtcgaattc | agatcgtagtctaccagctc |
| 13 | tctgcctctttgcgggaaac | cacggacatgggtgttgaat |
| 14 | ttcttggtttctggcagttc | gacatgggggactgaacatc |
| 15 | ctttttcacttccagtgagt | atgacaccgagaatggagcg |
| 16 | taataaggggcatcctggtc | cacgctgatatcactggact |
| 17 | cgtagtgactgcactgttgc | cggaagctccttgactcttc |
| 18 | tggtgggcatcgtagatgtg | tactgaaccacttccaggaa |
| 19 | cgcttgctggaaagggacat | atattccagaatgtccatct |
| 20 | tcaatgtcctcgattccagc | cgatgtttcttctcgctctc |
| 21 | caggttctccttgtcattgg | ttctgtctgcgcatatagag |
| 22 | cgtttacatattcggagacc | agcgcatattgtggctaatg |
| 23 | tccacctgatacaagtagtc | accagccaatcaataaggat |
| 24 | caggtgatccttgtgaatgg | cagtttgtactcctcggaaa |
| 25 | caatcgatcagcacggctcg | aagaccgagagatagagcgt |
| 26 | actgcaggtggacttcgttg | ccgccatttggctcaaaaat |
| 27 | aaggtctctgcagccagatg | agctgtaacttggagcgcac |
| 28 | gtagcgatcaatgatagcca | caatatacatagctgccgtg |
| 29 | attgcaagtacgtgcgtttg | gggatagatttcctcgtatt |
| 30 | gctatgaagagtgctgtcac | gaagacaaactcaccgacct |
| 31 | ggaacagctcctcgtacttg | cttggtgtaactgtcgtcgg |
| 32 | gacgaaatctccgattgccg | ttcaagataacctgctccat |
| 33 | tgtaggtgtcgtccgtgatg | tgcacagatcgaaggagaga |
| 34 | ttgaagatttgcagctccat | atgaagacgtaagcagtcgg |
| 35 | cgcgacagattacagtcgat | aggcatgtcgcaaagtacag |
| 36 | aaggaagtgaatcggcagcg | tacagcgtcatgtacttcag |
| 37 | gtacttggacatcgtatggt | ccatgagcgacaactcggaa |
| 38 | ccacggaagctaactcgatg | tattgcaagtacgtttctcc |
| 39 | ctgtaagtggccatttcgta | cgactgaagcggatgacata |
| 40 | cgacaggaacagtgaggcag | acatctccatgcccaaaatg |
| 41 | tggtttccattgagcaagtg | agcttgtaagtcgtgatctc |
| 42 | acggtcgttgaatcctgtac | gcaacacaaccgtttttagg |
| 43 | gagtatcgcgagtagaaggt | gttttgtgggtgtgacacag |
| 44 | cgggtaatcggacgcaagtg | tattgtacttttcccgcatg |
| 45 | tggtacttgttgtagatggc | catagccaccttcttgtaag |
| 46 | cgatcttctggaacttgctg | ttgctcatctcaacggattc |
| 47 | cacaatcgagtccatcagcg | cacagagctgatcgaagtcg |
| 48 | | tctttctgcttgcaattgta |
