## Supplemental Table S2 for "Post-transcriptional regulation of *cyclin A* and *cyclin B* mRNAs is mediated by Bruno 1 and Cup, and further fine-tuned within P-bodies"

**Table 2. Primer sequences used in RT-qPCR experiments**

| | |
|---|---|
| CycA FWD | 'GTCACACCCACAAAACGGCA' |
| CycA REV | 'GCTGCTGGTGCTCATCCTCT' |
| CycB FWD | 'CGAGGACGAGCACCATACGA' |
| CycB REV | 'AGTGCAGCGACAGGAACAGT' |
